# MPGEM: A harmonized and transcriptome-complete resource for large-scale reuse of legacy human microarray data

**DOI:** 10.64898/2026.08.20.746052

**Authors:** Shruti Gupta, Ajay Kumar Verma, Sougata Jana, Shandar Ahmad

**Author notes:** Equal contribution of authors (Shruti Gupta, Ajay Kumar Verma).

## Abstract

**Background:** Legacy microarray datasets provide an extensive record of human transcriptomic biology, but their reuse is constrained by differences in platform design, preprocessing, measurement scale, and gene coverage. Platforms measuring only subsets of genes cannot readily be integrated with higher-coverage platforms, limiting large-scale analysis and computational modeling.

**Results:** We developed Multi-Platform Gene Expression Matrix (MPGEM), a computational framework and resource for harmonizing and completing gene-expression profiles across heterogeneous microarray platforms. MPGEM uses a Reference Quantile Distribution (RQD) and generalized Reference Subset Quantile Distribution (RSQD) framework to transform profiles with different gene coverage onto a common quantitative scale. The MPGEM Engine, a multilayer perceptron, predicts expression of unmeasured genes from genes shared across platforms. Applied to Affymetrix GPL570, GPL571, and GPL96, MPGEM uses GPL570 as a 19,320- gene reference space comprising 12,712 predictor and 6,608 target genes. The resulting resource contains 207,135 human gene-expression profiles across 19,320 genes. Evaluation using masked GPL570 profiles yielded mean sample-wise Pearson and Spearman correlations of 0.944 and 0.939, respectively, and mean gene-wise correlations of 0.830 and 0.825. The lowest-performing 5% of target genes achieved a mean Pearson correlation of 0.683. MPGEM showed comparable or higher predictive performance than baseline mean imputation and K-nearest-neighbor approaches.

**Conclusions:** MPGEM transforms heterogeneous, partially measured legacy microarray profiles into a harmonized, transcriptome-complete representation, facilitating their reuse for large-scale transcriptomic analysis, biomarker discovery, systems biology, and machine learning. The framework, trained models, and expression resource are provided as open-source resources.

## 1 Introduction

Gene expression (GE) underlies virtually every biological process, and its dynamic regulation across tissues, developmental stages, diseases, and environmental perturbations has been a central focus of molecular biology for decades (Schena et al., 1995; Lockhart et al., 1996). Large-scale transcriptomic profiling has enabled numerous discoveries in disease mechanisms, biomarker identification, drug response, and systems biology (Golub et al., 1999; Van De Vijver et al., 2002; Van ’T Veer et al., 2002; Lamb et al., 2006). To facilitate data sharing and reuse, transcriptomic experiments have been systematically archived in public repositories, among which the Gene Expression Omnibus (GEO) has become the largest and most widely used (Edgar, 2002; Clough et al., 2024). GEO contains hundreds of thousands of human gene expression profiles generated under diverse biological conditions, representing an unparalleled record of transcriptional responses across diseases, drug perturbations, developmental stages, and cellular states (Clough et al., 2024). Collectively, these datasets constitute an invaluable resource for large-scale biological discovery, enabling questions to be addressed that extend well beyond the scope of individual studies through integrative, complex, and meta-analysis(Rung and Brazma, 2013).

Although RNA sequencing has largely replaced microarrays as the preferred technology for transcriptome profiling (Wang et al., 2009; McGettigan, 2013), legacy microarray datasets remain scientifically valuable. First, the sheer volume of archived microarray experiments makes regeneration of all recorded conditions on an NGS platform economically impractical. Second, many studies represent unique patient cohorts, biological samples, treatment regimens, or experimental conditions that cannot readily be reproduced. Third, several influential secondary transcriptomic resources and computational frameworks have remained rooted in microarray measurements. For example, the original version of Connectivity Map (Lamb et al., 2006) was constructed entirely from Affymetrix microarray profiles, while the subsequent L1000 platform derived its landmark gene selection and inference models from large collections of microarray data from GEO. Similarly, resources such as Chemical Effects in Biological Systems (CEBS) (Waters et al., 2007), DrugMatrix (Ganter et al., 2006), TG-GATEs (Igarashi et al., 2015), and TSUNAMI (Huang et al., 2021) have demonstrated the power of large-scale expression profiling for functional genomics, toxicogenomics, and drug discovery, mostly using microarray data (Cooper et al., 2007). Likewise, numerous diagnostic, prognostic, and therapeutic biomarkers have been derived from microarray-based expression signatures. The tools linking microarray and RNASeq data have also been reported, among which (Su et al., 2014; Korir et al., 2015) have shown that machine learning models trained on microarray data can successfully generalize to RNA sequencing datasets. These studies collectively demonstrate that legacy transcriptomic data remain highly informative and continue to support biological discovery in the era of next-generation sequencing directly or indirectly.

Despite these success stories, a collective analysis of microarray experiments remains challenging because datasets generated on different platforms are not directly comparable (Walsh et al., 2015). Each microarray “platform” employs a distinct probe design, resulting in differences in probe-to-gene mapping, gene coverage, and quantitative measurement scales (Ramasamy et al., 2008; Allen et al., 2012). Consequently, expression profiles generated on different platforms often contain different feature sets, different numerical distributions, and varying levels of gene-level coverage. Integrating such heterogeneous datasets typically requires extensive preprocessing, probe remapping, normalization, and removal of platform-specific biases (Shabalin et al., 2008). Furthermore, because only a subset of genes, which are shared across platforms, can usually be retained, a substantial fraction of experimentally measured information is discarded during cross-platform integration. This loss of information limits both statistical power and biological interpretation when combining data from multiple studies.

Numerous computational approaches have attempted to address these challenges. Probe-level annotation methods have improved probe-to-gene mapping, while normalization methods such as quantile normalization and batch-effect correction algorithms have facilitated integration of studies generated under similar experimental conditions but quantified differently (Bolstad et al., 2003; Johnson et al., 2007). More recently, methods such as Shambhala (Borisov et al., 2019, 2022) introduced the concept of normalizing expression profiles against an external reference distribution rather than one derived from the datasets themselves. Although these approaches significantly improve comparability, they generally assume a common feature space and therefore cannot reconcile experiments generated on platforms with unequal and non-identical gene coverage. To the best of our knowledge, existing frameworks do not simultaneously address cross-platform normalization and transcriptome completion across heterogeneous microarray platforms within a unified resource.

These limitations have become increasingly important with the emergence of artificial intelligence (AI) in biomedical research. Modern machine learning and deep learning algorithms require uniformly processed datasets with consistent feature representations across samples. Large-scale AI models cannot be effectively trained on heterogeneous expression matrices that differ in measurement scales, gene coverage, and missing features. Thus, the absence of a harmonized, gene-complete transcriptomic resource has become a major bottleneck not only for transcriptomic meta-analysis but also for developing foundation-scale AI models capable of learning generalizable biological representations from public gene expression data.

Multiple efforts have been made in the past to harmonize microarray data sets, which range from normalization methods to probe set mapping strategies. The widely used quantile normalization is not reusable on new samples, and for this purpose, two methods have been noteworthy. First, the fRMA method proposed many years ago uses a frozen reference for normalization to improve the reusability of references derived from larger sample pools to normalize new profiles (McCall et al., 2010). Along those lines, more recently, the Shambhala (Borisov et al., 2019, 2022) method has further refined this approach. However, most of these reference-based approaches are meant for identical gene set profile normalization and would not be usable if a new experiment does not contain all the genes included in the reference set. This is a key limitation of current reference-based reusable normalization strategies.

Attempts to provide a unified resource from multiple cross-platform experiments have also been around for a while. Among them, *refine.bio* (Greene et al., 2026) is a uniformly processed, cross-platform harmonized compendium of GEO/ArrayExpress/SRA, including microarray platforms, with quantile normalization to reference distributions. Similarly, *Gemma* (Zoubarev et al., 2012) is a curated reprocessed GEO microarray data at scale. However, all these resources integrated expression profiles within the scope of the originally available gene set, without attempting to expand the smaller sets to infer the large number of missing genes in some of the platforms,

The closest effort in this direction was made by D-GEX (Chen et al., 2016), which reported a deep learning method for inference of ∼9,500 target genes from 978 landmark genes. However, both the target and predictor genes in this model were taken from the same platform (GPL96), and many genes from even this platform were not included in the model. The usability of D-GEX for cross-platform prediction and for the many other genes from the same platform is unclear, and therefore, the model is not directly usable for completing the large number of transcriptome profiles in GEO.

To address these challenges, here we report the development of the **Multi-Platform Gene Expression Matrix (MPGEM)**, a unified human microarray resource that harmonizes expression profiles across major GEO microarray platforms and reconstructs missing gene expression values through artificial intelligence-based predictive modelling. The overall framework for achieving this complex goal consists of three complementary components. First, we develop a platform-independent harmonization strategy based on a **Reference Quantile Distribution (RQD)-similar to fRMA and Shambhala-2** but expand it to a more generalized framework, **Reference Subset Quantile Distribution (RSQD)**. The new framework enables consistent normalization of expression profiles generated from high gene coverage samples as well as their arbitrary subsets. Second, we develop the **MPGEM Engine**, a multilayer perceptron (MLP)-based model trained on high-coverage expression profiles to accurately predict expression values for genes absent from lower-density microarray platforms. Finally, we integrate these methods into a unified transcriptomic resource comprising harmonized and expanded gene expression profiles together with software tools for processing newly generated or legacy microarray datasets using the same framework.

Our current MPGEM release contains 207,135 profiles uniformly represented across 19,320 genes. The accompanying MPGEM Engine reconstructs missing expression values with a mean Pearson correlation of approximately 0.944, providing accurate transcriptome completion across heterogeneous platforms. By combining robust cross-platform harmonization with AI-based gene expression imputation, MPGEM establishes a comprehensive transcriptomic resource for large-scale meta-analysis, biomarker discovery, systems biology, and AI-driven functional genomics, while providing a scalable foundation for future integration with RNA sequencing and other transcriptomic technologies.

## 2 Material and Methods

The MPGEM pipeline proceeds in four scalable stages, shown in Figure 1. As shown in this Figure, gene expression data from three human microarray platforms was first collected from GEO. Choice of included platforms was made manually based on their numerosity, gene set coverage, and overlap with the CMAP and L1000 gene sets (explained in section 2.1). In the second stage, the probe-level expression data in the consolidated platform files were mapped to gene identifiers, cleaned, and assembled into a composite raw expression matrix per platform, the pre-MPGEM matrices (see section 2.2). In the third stage, these matrices were harmonized into a single common scale using a rigorously computed Reference Quantile Distribution and its subset-adapted generalization, both of which were introduced in this work (see section 2.3). In the fourth stage, the genes absent from the two smaller platforms were extrapolated using an MPGEM Engine, an MLP-based model trained on the harmonized GPL570 data, producing the complete MPGEM matrix (see section 2.4). The engine is evaluated and benchmarked against baseline methods in the corresponding Results sections. Throughout this study, GPL570 serves as the reference platform, since the gene sets of the other two platforms are confirmed to be a subset of these genes in this platform.

**Figure 1:**
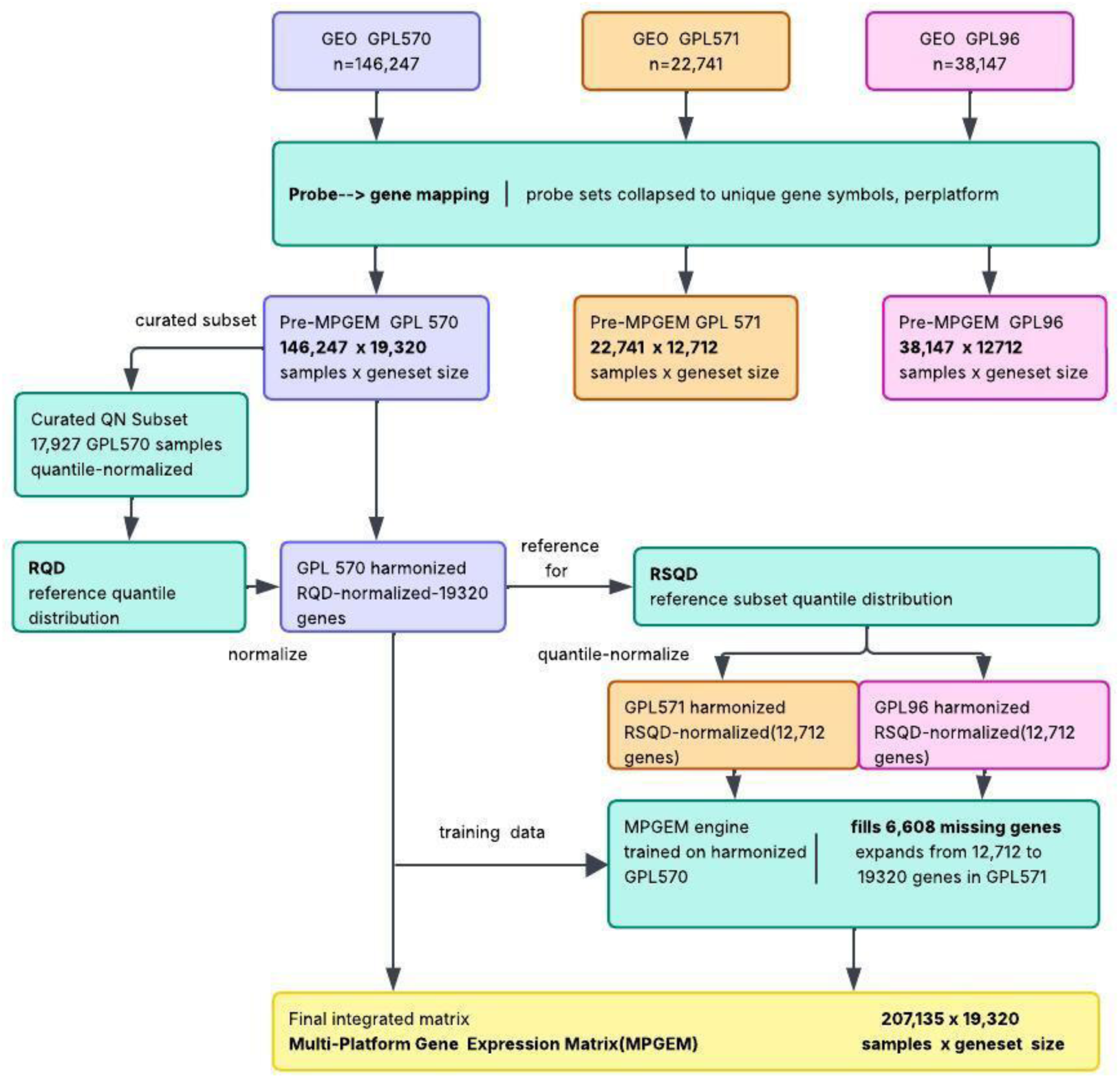
Overall pipeline followed in this work from pre-processing to training MPGEM matrix.

### 2.1 Microarray chip selection from the GEO Database

Millions of gene expression profiles are available in GEO and ArrayExpress (Athar et al., 2019), collected using different technologies and platforms for genomes representing diverse species. In this work, we focused on data from Homo Sapiens only and selected three oligonucleotide microarray chips in GEO with the highest abundance of samples, namely GPL570, GPL571, and GPL96. Apart from their abundance in GEO, these platforms were also important because of their coverage of the genes included in the CMAP and L1000 gene sets, a drug discovery platform derived for identifying minimal gene sets for expression prediction (Subramanian et al., 2017). Relating to CMAP/L1000 is helpful in utilizing the gene sets already grouped in terms of core and inferred genes in that study. Each platform’s data contains at least one probe set for each of the 978 genes included in the L1000 landmark sets. It may be recalled that the samples from GPL96 were the primary source for L1000 in (Subramanian et al., 2017), which is one of the smaller platforms selected here.

### 2.2 Construction of pre-MPGEM raw matrices

As stated above, each GEO platform record, identified by a GPL prefix, defines the design of a microarray and indexes every sample deposited from experiments performed on it. We retrieved all samples (GSM records) indexed under selected platforms using the compressed “Platform files” from GEO (https://www.ncbi.nlm.nih.gov/geo/; downloaded in September 2023, but the data have largely remained unchanged since). Extraction of sample-wise GE data from the platform file was performed using an in-house Python code. For microarray platforms, these data are recorded at the probe level in these extracted profiles. We converted these probe-level data to gene level (see 2.2.1) and created a raw matrix in which rows are GSM sample IDs and columns are gene names. Profiles were cleaned for anomalies, within-platform missing values, and mapping ambiguities (see 2.2.1, 2.2.2), giving one raw matrix per platform.

#### 2.2.1 Translating probe information to gene identifiers

Probe-to-gene assignments were taken from the annotation files provided by Affymetrix, the most recent release of which is version 36 (2016) (https://www.thermofisher.com/bd/en/home/life-science/microarray-analysis/microarray-data-analysis/genechip-array-annotation-files.html). Because gene models have been revised since then, every assignment was manually rechecked against the NCBI Gene database before use. We retained only probe sets mapping uniquely to a single gene, without resolving isoforms. Probe sets with no gene mapping, with multiple gene mappings, or whose target gene has since been retired in NCBI Gene were excluded. To convert the expression levels of multiple probes corresponding to the same single gene, we took the maximum value across all probes for that gene within each sample (Miller et al., 2011; Subramanian et al., 2017). Even though it implies that different probes may be selected for a given gene in different samples, it avoids diluting a true signal with probe sets that cross-hybridize poorly or target degraded transcript regions, and it is applied independently within each sample, so no cross-sample information leaks into the mapping (Eq. 1):

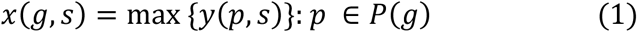

where *y*(*p*, *s*) is the expression value of the probe set *p* in sample *s*, *P*(*g*) is the set of probe sets mapping uniquely to gene *g*, and *x*(*g*, *s*) is the gene-level expression of *g* in *s*.

#### 2.2.2 Addressing missing probe-level values

Some samples across all three platforms contained one or more missing probe values. These were substituted with the minimum value from all probes of all genes observed within that sample (Eq. 2). A missing value most commonly reflects a probe whose signal fell below reliable detection, so substituting the sample minimum treats the transcript as present at or below the detection floor rather than absent at random:

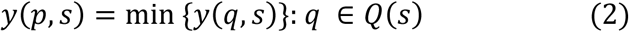

for all *p* missing in *s*, where *Q*(*s*) is the set of probe sets with recorded intensities in sample *s*.

### 2.3 Processing of pre-MPGEM gene expression matrices

The outcome of the previous step is a pre-MPGEM matrix, a gene-mapped filtered set of gene expression values from the three platforms as described above. The component pre-MPGEM matrices must be harmonized individually, integrated onto a common scale, and then extrapolated into a single multi-platform matrix, the final MPGEM. Harmonization is performed by assigning an identical distribution to each sample in a manner of quantile normalization. In order to preserve the quantile-normalization distribution, we aimed to create a robust universal distribution, which can be reused without going through the quantile normalization steps from scratch. We set certain quality requirements for constructing this. First, the reference distribution must be built only from samples already quantile-normalized and verified to match that label, so that it represents averages over comparable scales. Second, the reference had to be reusable on new samples, not in the current dataset, so the resource remains usable as new profiles are deposited. Third, it had to apply to samples covering only a subset of the gene set. Ordinary quantile normalization cannot do this because it would re-estimate the target distribution from the smaller gene set alone, thereby losing the ranks of the shared genes within the full transcriptome. We therefore developed a procedure that applies the reference built on the full gene set to samples with fewer genes, assuming the incoming gene set is a subset of the reference. For the full-scale reference distribution usable for new samples without recalculating the distribution, we created a robust approach of deriving a reference distribution from high-quality profiles defined as follows. Thus, the harmonization requirements were fulfilled as follows.

#### 2.3.1 Selection of samples for Reference Quantile Distribution (RQD)

First step in a typical quantile normalization method for gene expression involves averaging of quantitative gene expression values at specific ranked positions within each sample profile being normalized. In a heterogeneously normalized data set, such an approach may end up taking averages of unrelated distributions. In other words, a reference distribution without segregating similarly scaled studies may end up inadvertently computing averages of similarly ranked GE values originally on very different scales. To overcome this problem, we constructed a universally reusable Reference Quantile Distribution (RQD) from a curated subset of GPL570 samples. This step of the method is similar in principle to the idea of a fixed external reference defined recently (Borisov et al., 2019, 2022). We have used a much more rigorous approach here and go beyond the original scope in selecting reference distribution and enabling its adaptation to a smaller gene subset, for which full-scale ranks in the RQD could not be computed. We derived our RQD from samples already quantile-normalized and on a consistent scale. Metadata labels of quantile normalization suggested that they cannot be taken at face value, i.e., filtering samples on the “quantile” keyword was necessary but not sufficient to select similarly normalized values, so we manually verified that each sample’s expression values were consistent with its stated scale.

After careful curation of a high-quality set of log2 quantile-normalized GPL570 samples, we constructed the Reference Quantile Distribution (RQD) as follows. Within each sample, gene expression values were sorted in ascending order and assigned ranks 1 to N, where N = 19,320 is the number of genes retained after probe-to-gene mapping. At each rank, we took the mean expression across all selected samples (Eq. 3). The resulting vector of one mean per rank is the RQD, a consensus expression profile drawn from the curated quantile-normalized samples:

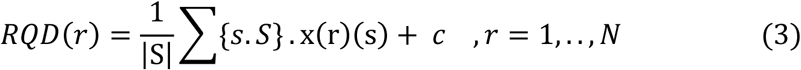

where, *S* is the curated sample set, x(r)(s) is the r ^th^ smallest value in sample s, *N* is the gene set size (=19,320 here) and *c* is an offset value added to all expression values to rule out negative values, which cannot be handled by our activation function (*c* = 2.0 here).

Because an offset is applied to every profile, it does not significantly affect any rank- or correlation-based comparison. Supplementary Table ST3 shows the distribution of the RQD. As an alternative to the mean, we also built candidate RQDs from trimmed ranges (percentiles 1–99, 0.1–99.9, 0.01–99.99) to check whether extreme ranks distorted the reference; the distributions were essentially unchanged, so the mean-based RQD was retained. The RQD was then used to normalize every sample in the pre-MPGEM GPL570 matrix: expression values were rank-ordered and replaced with the corresponding RQD entries (Eq. 4). Tied values within a sample were assigned averaged ranks, truncated to an integer index, so all genes in a tie group map to the same RQD entry. This is more stable and less sensitive to study-specific artifacts, and provides a fixed target for normalizing new profiles:

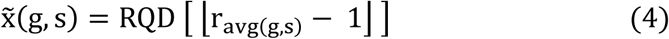

where r_avg(g,s)_ is the average (1-indexed) rank of gene g in sample s under tied ranking, [.] is the truncation, and RQD [.] is 0-indexed.

#### 2.3.2 Subset Quantile Distribution (RS-QD) for universal adaptation of RQD

We developed the following strategy for adopting RQD for gene sets with a smaller coverage. We redefined the reference distributions for subsets by first extracting the values of the genes shared with the smaller profiles (GPL571 and GPL96 here), sorted them within each sample of the larger set data (RQD-normalized pre-MPGEM-570 data), assigned ranks 1 to M (M = 12,712), and took the rank-wise mean across samples (Eq. 5). The resulting vector is the Reference Subset Quantile Distribution (RSQD): the consensus profile of the shared gene set, carrying forward the rank relationships established in the full-length RQD rather than being re-estimated on the smaller platforms.

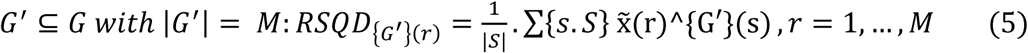

where x̃(r)^{G′}(s)is the r ^th^ smallest RQD-normalized value among the genes of *G*^′^ in sample s, and *M* = size of the subset (*M* =12,712, for GPL96 and GPL571)

The RSQD was then used to normalize each sample in the pre-MPGEM GPL571 and GPL96 matrices (Eq. 6). Because both platforms map to the same 12,712-gene set, a single RSQD was used for both.

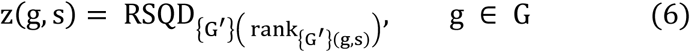

where rank_{G_′_}(g,s)_ is the ascending rank of gene g among the G^′^ values of sample s.

Definition of RSQD is shown here for GPL571 and GPL96, but is scalable to a GE profile for any arbitrary gene subset of GPL570, which is treated as a global superset of low coverage platforms. The overall normalization strategy used is called the Reference Subset Quantile Distribution (RSQD) method in which for any new GE profile, an RSQD is generated based on its gene coverage and used to normalize it. The RQD is the special case where the profile covers the full 19,320-gene, and hence the RSQD reduces to the full-scale RQD. The defining property of the method is that the reference is always derived from the RQD-normalized GPL570 matrix, never re-estimated on the smaller platform. This keeps the shared genes’ ranks consistent with their positions in the full transcriptome and places all platforms and new partial profiles on a common scale. The harmonized matrix is assembled by normalizing each pre-MPGEM matrix (GPL570, GPL571, GPL96) to its corresponding reference and then stacking the results. At this stage, the GPL571 and GPL96 profiles carry values for only 12,712 of the 19,320 genes in GPL570; extrapolation of the remaining 6,608, which completes the MPGEM, is described in 2.4. The pre-MPGEM now has comparable expression values in each column after this normalization and is called pre-MPGEM-n for reference. This matrix has missing values in columns of lower coverage platforms from which platform-wise subsets of this matrix are extracted as pre-MPGEM-570-n, pre-MPGEM-96-n, and pre-MPGEM-571-n without any missing values each.

### 2.4 MPGEM imputation engine

To obtain a fully populated cross-platform gene expression matrix from sparsely populated pre-MPGEM-n, we developed an imputation engine based on a multi-layer perceptron (MLP)- the MPGEM-engine, which predicts the gene expression values absent in GPL96 and GPL571 from the genes they share with GPL570. The engine is trained on pre-MPGEM-570-n data by masking these genes to be extrapolated and then reconstructing them from the remaining genes. Once a predictive model is trained, it is applied to pre-MPGEM-96-n and pre-MPGEM-571-n to impute their missing gene block. The predicted values are combined with the measured values to yield the complete MPGEM: 207,135 samples across all 19,320 genes. Overall architecture of MPGEM engine and the number of genes to predict is illustrated in Figure 2(a).

**Figure 2:**
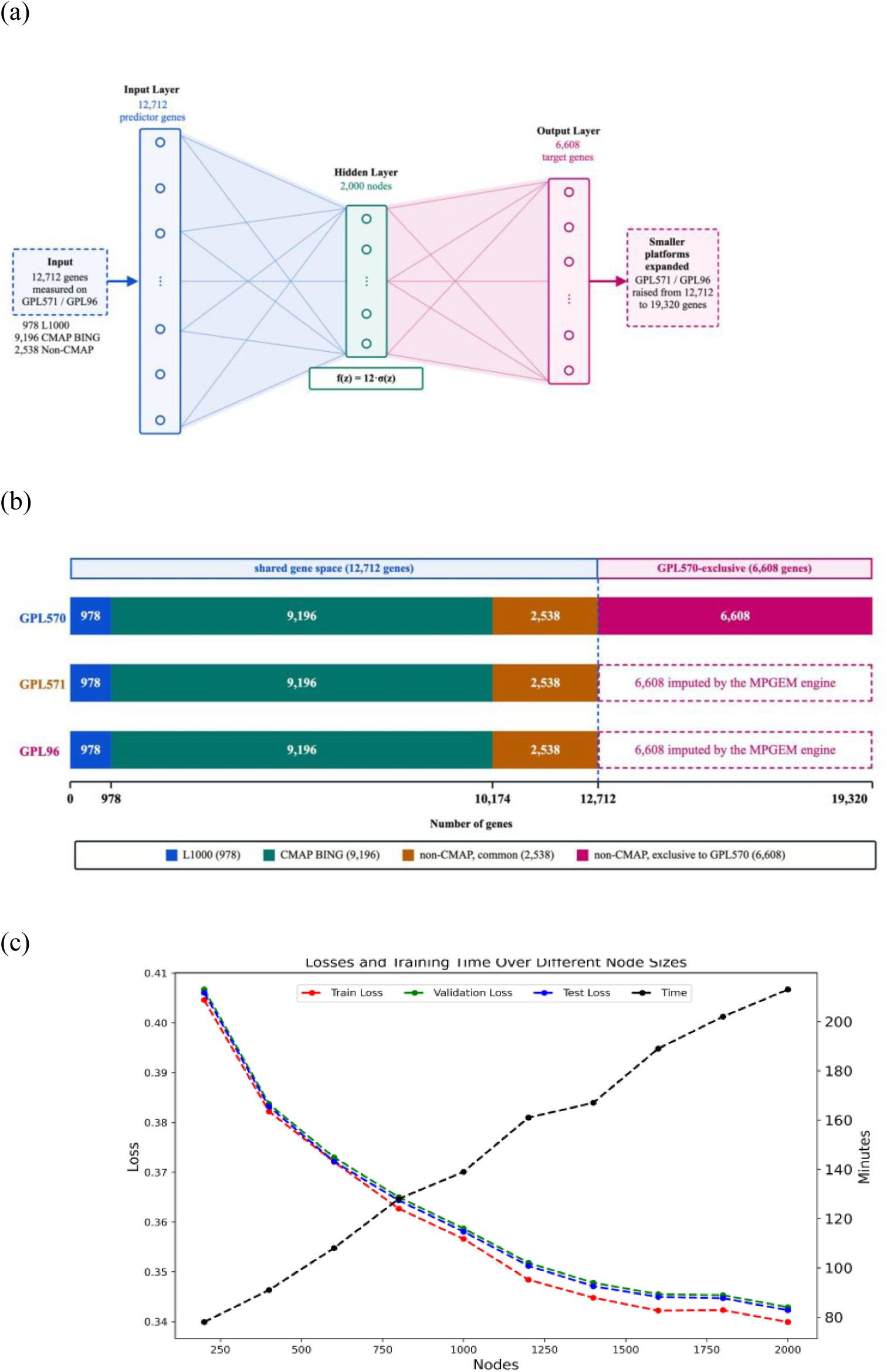
Training strategy for MPGEM (a) MPGEM-engine design for training and (b) Gene set coverage in selected GPL-prefixed platform data. All the genes included in GPL96 and GPL571 genes (including L1000, BING, and non-CMAP genes) form a subset of GPL570 genes, making the latter usable to train MPGEM engine (c) (b) Training, validation, and test loss (left axis) and training time (right axis) versus hidden-layer size. Loss flattens beyond ∼1,500 nodes while training time rises approximately linearly; the three loss curves coincide, indicating no overfitting.

#### 2.4.1 Predictor and target gene definition

The genes retained on GPL570 after probe-to-gene mapping were partitioned by their availability on the smaller platforms, as illustrated in Figure 2(b). The 12,712 genes shared across all three platforms formed the predictor set, and the 6,608 genes present on GPL570 but absent from GPL96 and GPL571 formed the target set, together accounting for all 19,320 GPL570 genes. For reference, the predictor set comprises the 978 L1000 landmark genes, the 9,196 “Best Inferred Genes” (BING) defined by the CMAP project (Subramanian et al., 2017), and 2,538 shared genes outside the CMAP designation, which have been addressed for the first time in this work. The engine learns the predictor-to-target mapping from pre-MPGEM-570-n, where both groups are measured, and applies it to pre-MPGEM-96-n and pre-MPGEM-571-n, where only the 12,712 predictors are available.

#### 2.4.2 MLP architecture of MPGEM engine

The MPGEM-engine is a feed-forward multi-layer perceptron for multi-task regression, with one input layer, a single hidden layer, and one output layer. The input layer has 12,712 nodes, amounting to one per predictor gene; the output layer has 6,608 corresponding to as many target genes. Each hidden node computes a weighted sum of the preceding layer’s outputs plus a bias, and applies a non-linear activation (Eq. 7):

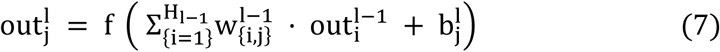

where 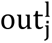 is the output of node j in layer l, 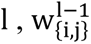 is the weight from node i in layer l − 1 to node j in layer l, 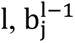 is the bias of node j in layer l − 1, H_l−1_is the number of nodes in layer l − 1 and f is the activation function used.

The hidden layer uses a scaled sigmoid activation (Eq. 8):

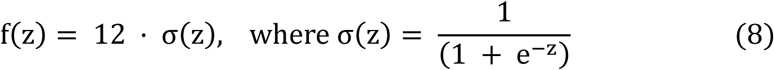

The factor of 12 matches the effective dynamic range of the RQD-normalized values, so hidden-layer outputs span the data range rather than being compressed into the unit interval of the standard sigmoid. The output layer uses a linear activation, appropriate for continuous regression. Weights and biases were optimized by minimizing the mean absolute error (MAE) between predicted and measured expression across all target genes (Eq. 9):

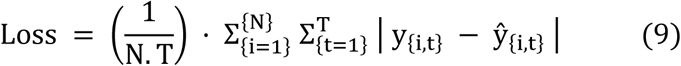

where N is the number of training samples, T is the gene set size (T =6608 here) and y_{i,t}_ is the measured value of target gene t in sample i, and ŷ_{i,t}_ the prediction.

#### 2.4.3 Regularization and weight initialization

Network weights were initialized by Glorot uniform initialization (Glorot and Bengio, 2010) to stabilize the variance of activations and gradients during training. We evaluated dropout during architecture selection but did not retain it, as it did not improve validation loss; the final engine was trained with a dropout rate of zero.

#### 2.4.4 Training procedure

The samples were partitioned at the sample level into 64% training, 16% validation, and 20% independent test sets. The training set was used to optimize model parameters, the validation set was used for architecture and hyperparameter selection, and the independent test set was used only for final performance evaluation. No sample was shared between the three subsets. Splitting was at the sample level, so no sample appeared in both training and evaluation. The learning rate was fixed at 1×10^-6^. Training continued until the maximum allocation of 10,000 epochs because the validation performance did not decline sufficiently to trigger the predefined early-stopping criterion.

#### 2.4.5 Architecture selection

The hidden-layer size controls the engine’s capacity and was chosen empirically. We trained the MPGEM-engine with hidden-layer sizes from 200 to 2,000 nodes and evaluated each on the validation set of pre-MPGEM-570-n. Training and validation loss fell together as hidden-layer size increased and flattened beyond ∼1,500 nodes, while training time rose approximately linearly (Figure 2(c)). The independent test set was not used for architecture selection. Because accuracy gains past ∼1,500 nodes were marginal and came at rising compute cost, we adopted 2,000 nodes as the plateau.

#### 2.4.6 Implementation

The MPGEM engine was implemented in Keras (Chollet and others, 2015) on the TensorFlow backend (Abadi and others, 2015) and trained on an Nvidia Quadro RTX 6000 GPU (24 GB memory, 4,608 CUDA cores).

### 2.5 Evaluation of imputation performance

The imputation engine must be evaluated on genes for which a true measured value exists, so that predictions can be checked. We therefore evaluated it on held-out GPL570 samples, where every gene is measured, by masking the target genes and comparing the engine’s predictions against their measured values.

#### 2.5.1 Masking design

The target genes are absent from GPL96 and GPL571, so no measured values exist there to check a prediction against. Evaluation was therefore done on GPL570, where all 19,320 genes are measured. Using the test split of pre-MPGEM-570-n (2.4.4), we masked the 6,608 target genes in each test sample, leaving only the 12,712 predictors visible to the engine. The engine predicted the masked genes, and each prediction was compared with the measured value for that gene in that sample. Because a masked test sample reproduces exactly the predictive features of a GPL96 or GPL571 sample, with only the 12,712 shared genes available, this measures performance on the real imputation task rather than a proxy.

#### 2.5.2 Performance metric

Agreement between predicted and measured expression was quantified by parametric and non-parametric coefficients of correlation namely the Pearson and Spearman correlation coefficients, each computed in two directions. In the sample-wise direction, the correlation was computed across the 6,608 target genes within each test sample and then averaged over samples; this measures how well each sample’s target profile is reconstructed. In the gene-wise direction, the correlation was computed across all test samples for each target gene and then averaged over the 6,608 genes; this **measures how well the engine reproduces each gene’s variation across samples. Pearson correlation is** defined as (Eq 10)

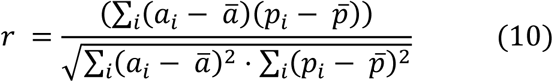

where *a_i_* and *p_i_* are the measured and predicted values, and *a̅*, *p̅* their means over the set being correlated.

#### 2.5.3 Performance comparisons

Because MPGEM and KNN predictions (see Results) were evaluated on the same test samples, differences in sample-wise performance were assessed using paired t-tests. For gene-wise comparisons, the same pairing principle was applied across the target genes.

## 3 Results

### 3.1 Data download summary and pre-MPGEM

Number of samples and probe count in the platform files GPL570, GPL96, and GPL571, downloaded from NCBI is provided in Table 1, which are utilized to construct the pre-MPGEM data. As noted, both GPL571 and GPL96 contain near-identical probe sets (22,277 and 22,283, respectively), both contained within the larger probe set of GPL570. After probe-to-gene mapping, the two resolve to an identical gene set of 12,712 genes, a strict subset of the 19,320 genes, available from GPL570, not used earlier in CMAP.

**Table 1:** Prior-normalization breakdown of the GPL570 samples, and the subset curated for RQD construction.

| Details | pre-MPGEM<br>GPL570 |
| --- | --- |
| Quantile normalized (Total) | 17,927 |
| at raw scale, no log | 10,298 |
| at log2 scale | 7,175 |
| at log10 scale | 454 |
| Non-quantile-normalized,<br>ambiguous scaled and others | 128,320 |
| % samples for RQD construction | ~12.3% |

### 3.2 Challenge of harmonization

Supplementary tables ST1 and ST2 show that the basic statistics of expression levels in all the samples, even from the same platform, are widely different, clearly showing the challenge of bringing **data to the same level of quantification. After performing the harmonization task on this platform** alone, all profiles are assigned the same distribution, which is normal-like for GPL570 and produces a slightly bimodal but near-normal distribution for the RQD. Detailed results on the subsequent steps are provided below.

### 3.3 Statistical distribution of original normalization methods in pre-MPGEM

As stated, pre-MPGEM is a sparse matrix consisting of data from three selected platforms at different levels of coverage. Its subsets are fully populated matrices of the size of the gene set representing these platforms, but are not uniformly normalized in the raw data files. In order to define a global reference distribution for RQD (see Methods), we examined the pre-existing normalization protocols in the source data files. Table 2 describes the observed number of normalization methods in the source files from pre-MPGEM-GPL570. As seen from this table, only ∼12% (17927 samples) were originally quantile-normalized on a log2 scale, which nonetheless present a robust set for reference quantification of GE data. Our proposed RQD is derived from these coherently quantified expression profiles.

**Table 2:** Details of microarray chip and associated raw experimental data downloaded from GEO.

| GEO ID |  | GPL570 | GPL571 | GPL96 |
| --- | --- | --- | --- | --- |
| Description |  | [HG-U133_Plus_2]<br>Affymetrix<br>Human Genome<br>U133 Plus 2.0 Array | [HG-U133A_2]<br>Affymetrix<br>Human Genome<br>U133A 2.0 Array | [HG-U133A]<br>Affymetrix<br>Human Genome<br>U133A Array |
| Raw<br>download<br>statistics | Total probe sets in array | 54,675 | 22,277 | 22,283 |
|  | No. of available samples | 177,329 | 24,479 | 43,657 |
|  | Latest data added in GEO | 2020 | 2018 | 2021 |
| Pre-MPGEM<br>contents | No. of samples retained | 146,247 | 22,741 | 38,147 |
|  | No. of genes annotated | 19,320 | 12,712 | 12,712 |
|  | No. probe sets mapped | 41,115 | 20,225 | 20,225 |
|  | No. of L1000 genes | 978 | 978 | 978 |
|  | No. of CMAP genes | 9,196 | 9,196 | 9,196 |
|  | No. of non-CMAP genes | 9,146 | 2,538 | 2,538 |
|  | No. of GPL570-exclusive genes | 6,608 | 0 | 0 |

Supplementary Table ST1 (a-d) shows how the basic statistics of mean, variance, maximum, and minimum expression values vary in the rows (samples) and columns (genes) of pre-MPGEM data. Supplementary Table ST2 shows some sampled distributions from these sets. Since some of these values are really outliers, we trimmed the data to show only the allowed ranges of percentiles in each plot so that a broad distribution can be visualized. We observed that the mean expression values of selected samples vary widely across samples and follow a multimodal distribution. When the trimming range was reduced to include more outliers, we observed that the mean values of small percentages (yet thousands of samples) were extremely high and suggested that the quantification of these samples was highly unusual, which could actually represent quantification errors. Multimodality of distributions into three or four distributions also suggested that these profiles formed certain groups, probably in terms of how the data were originally collected or processed. While sample-wise statistics showed these anomalies, they were not visible in gene-wise distributions, which appear to follow a simple negative exponential distribution. This suggested that gene-wise outlier distributions were mitigated by variations on both sides of the distribution. Since the critical property of interest from a gene’s perspective was to unambiguously assess its expression in a sample representing a biological context, the averaging effect on their expression values was not sufficient to use them without further processing.

Supplementary Table ST1 shows that the sample-wise variances of pre-MPGEM data are also widely different. Under these conditions, comparing gene expression values from one study to another may pose a huge statistical challenge, and thereby such distributions are not suitable for meta-analysis or cross-platform integration of biological knowledge. Maximum and minimum distribution patterns also support this argument. Minimum values for many samples run in the negative range, throwing up yet another question on the suitability of a standard quantile normalization-based harmonization of these data sets. In order to develop better insights into how the samples in pre-MPGEM differ from one another in terms of the entire distribution of their expression values, we plotted some representative profiles by selecting several samples randomly and then visually grouping them in terms of typical patterns. These are discussed in the next section.

To evaluate whether a standard quantile distribution on the overall pre-MPGEM is worthwhile, we looked at the typical distributions in these samples (rows of this matrix). Supplementary Table ST2 shows some of the histograms that were obtained when single sample expression profiles were analyzed. We observe that the scales and shapes of the gene expression value distributions widely varied. Some of these samples were highly kurtotic, bimodal, skewed, and followed non-normal distributions. Closely examining the meta-data, we found that the samples had been pre-normalized before submission to GEO and it was not straightforward to parse every single sample to know a-priori what normalization or scaling was already employed. For studies on small numbers of samples, such an annotation can be carried out manually but an automated method cannot produce exact answers. We, therefore, decided to collect samples that have a clear annotation of log2+quantile normalization in GEO and could be picked up with high confidence. Supplementary Table ST2 shows some of the samples picked up from such filters. We do observe a significantly more consistent distribution of expression values in this category of samples. Some variations do occur because the samples come from different batches or series and have different quantile distributions. Nonetheless, we feel that the samples collected from this filter can serve well to create a universal distribution of gene expression to be assigned directly to the ranked values of gene expression in any new sample profile. This observation and logic form the basis of our first level of the proposed quantile normalization-based harmonization method of this work.

### 3.4 Reference quantile distribution summary

Supplementary Table ST3 shows the final distribution generated in this way. Note that a constant value of 2.0 was added to all the first stage quantile values to avoid negative values in any of the gene expression positions in the matrix. We also tried another representative from the list of expression values on the n’th rank namely percentile values instead of average values. The distributions obtained by these metrics are shown in Supplementary Table ST3. However, we did not see a major change in terms of distribution patterns going from mean to these percentile values and therefore retained the standard average-based RQD as in Supplementary Table ST3. As seen from this supplementary table, the RQD so selected has a slightly bimodal pattern near its means, suggesting that the expression values fall into two groups, presumably into baseline unexpressed and second level expressed states. Finally, due to this bimodality and systematic ties in some locations, the final distributions in the normalized samples have a slight dip in frequency. However, the range of the mean values is really small as seen in Supplementary Table ST3. We therefore believe that the proposed RQD is a powerful reference distribution, retaining the properties of most real transcriptomes available in the public domain. The subset reference created for these two sets based on this approach is shown in Table 7 together with their distribution properties and gene-wise distributions achieved by this normalization. We observed from Supplementary Table ST3 that the subset reference extracted from RQD is similar to the GPL570 full length RQD- it is also bimodal as the former but the first peak is slightly lower than the second in the distribution function. The mean and variance distributions follow similar patterns. The true worth of these processed values can be assessed in their predictability and functional analysis that can be carried out over the integrated data as examined in the following sections.

### 3.5 MPGEM-engine and training results

MPGEM-engine hyperparameters were optimized by systematically increasing the size of the hidden layer in this model. Figure 3 shows the outcomes from the optimization process and the final predictive performance obtained in predicting the expression values of the target genes. Training and validation loss coincided throughout, with no apparent divergence indicative of substantial overfitting (Figure 3(a)). As stated in the Methods, it implies that the drop in validation data performance, which was used for early stopping did not occur and model continued to slowly learn until its maximum limit of 10,000. At the selected size, the final model was taken at 10,000 epochs. Figure 3(a) shows a typical learning curve. In the Methods section, we discussed (Figure 2(c)) that the amount of time taken for training the model for the same number of nodes increases roughly linearly in the range considered. Therefore, the prediction model for the application stage is the one trained using 2000 hidden nodes and 10,000 epochs. The performance levels for different number of nodes was tested on a smaller number of 5000 epochs (to save computing time) and their evolution with the increasing number of nodes are shown in Figure 3(c). During architecture optimization, models trained for 5,000 epochs achieved sample-wise Pearson correlations of approximately 0.91-0.93 and Spearman correlations of approximately 0.71-0.80, with performance plateauing beyond ∼1,500 hidden nodes. After selecting 2,000 hidden nodes, we trained the final model for 10,000 epochs. The final model achieved mean sample-wise Pearson and Spearman correlations of 0.944 and 0.939, respectively, and corresponding gene-wise correlations of 0.830 and 0.825 (Table 4).

**Figure 3:**
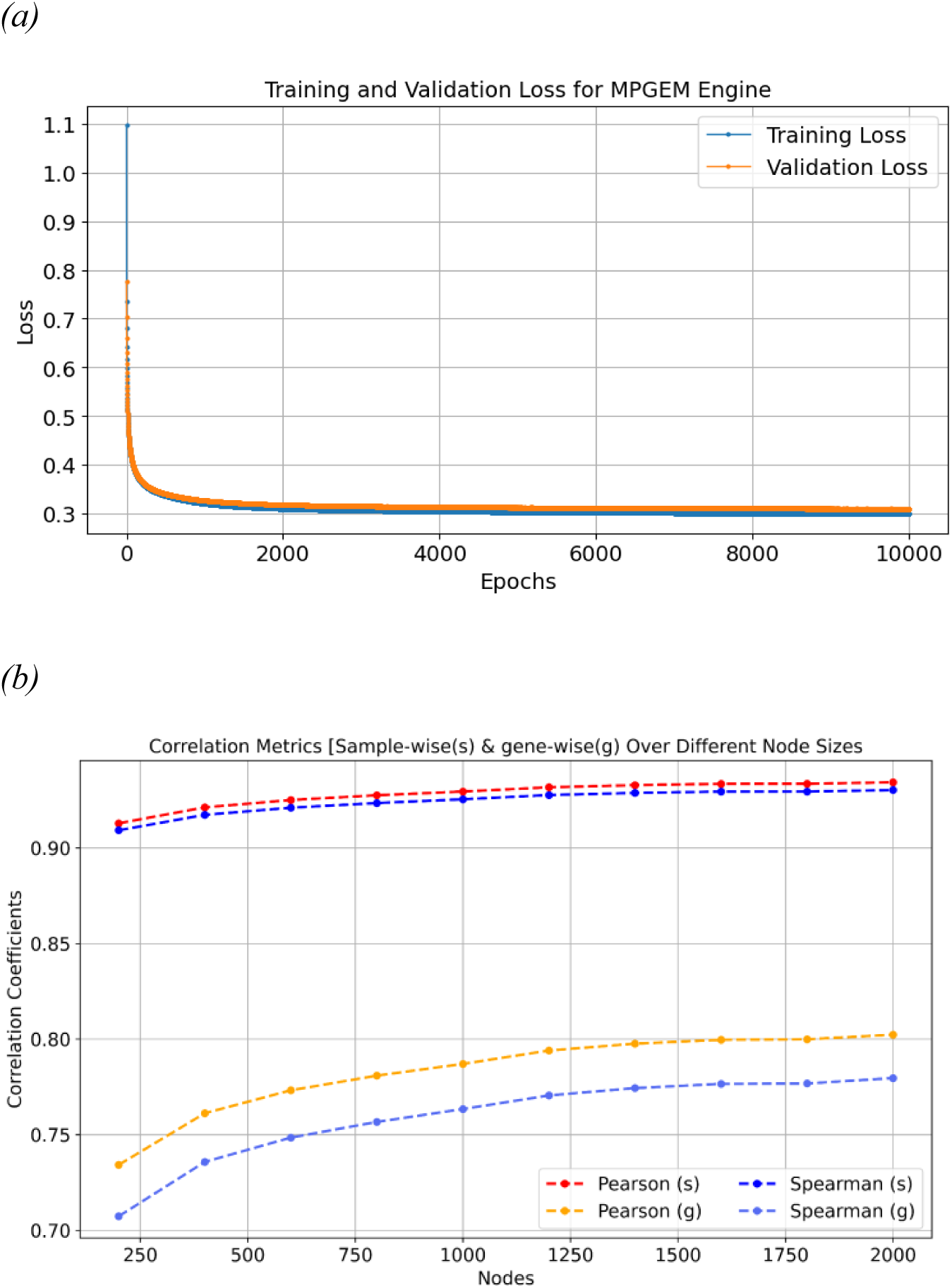
The training results of pre-MPGEM-n:GPL570 on different sizes of hidden layers (a) Training and validation loss (mean absolute error) for the MPGEM-engine over 5,000 epochs. The two curves coincide and continue to decline slowly without evidence of overfitting. Early stopping was defined based on a decline in validation performance, but this criterion was not reached during training. (b) Sample-wise (s) and gene-wise (g) Pearson and Spearman correlations between predicted and measured expression on masked GPL570 test samples, for hidden layers of 200–2,000 nodes obtained for 5000 epochs in order to estimate the impact of hidden layer size. Sample-wise correlations for this plot (∼0.91–0.93) exceed gene-wise correlations (∼0.71–0.80) and all curves plateau beyond ∼1,500 nodes. Further training for 10,000 epochs was carried out only for the finally selected (largest) network size (2,000 nodes) due to computational cost involved, results of which are slightly better than this plot (see Table 4).

#### 3.5.1 Gene-wise prediction performance

Although the overall mean sample-wise Pearson correlation was 0.944, it is useful to examine the distribution of prediction performance across individual genes. Figure 4(a) shows the histograms of Pearson and Spearman correlations as a metric of performance in predicting all the masked genes (those, which need to be imputed for GPL96 and GPL571). We observe that overall performances are robust across genes with the worst performing genes still showing good enough correlation coefficients of ∼0.6. Only a few hundred genes had correlations below 0.7, whereas a comparable number showed correlations of approximately 0.9 or higher.

**Figure 4:**
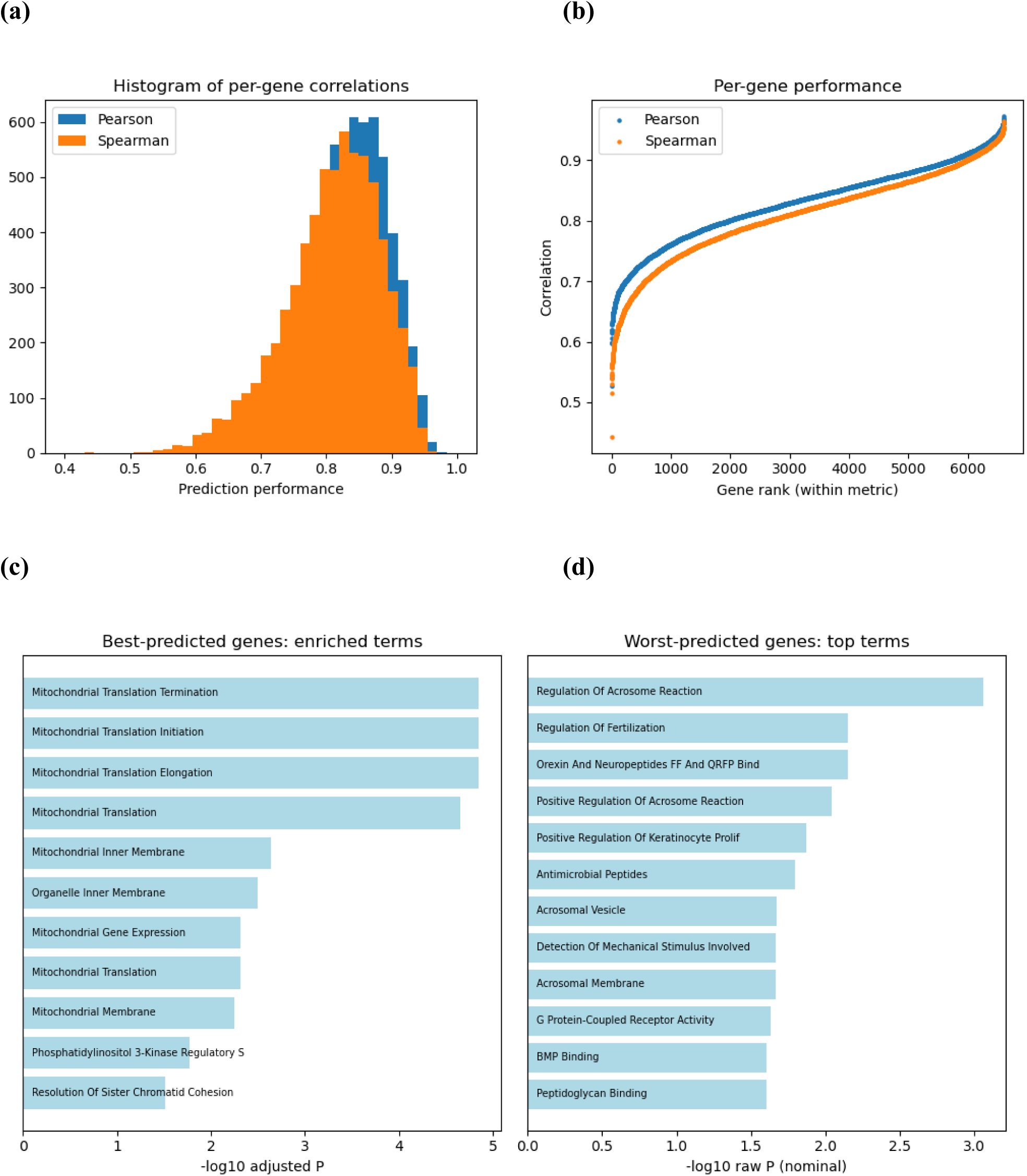
Gene-wise distribution of prediction power of MPGEM engine. (a) Overall distribution of correlation coefficient between predicted and target values, which shows good correlation for most of the genes with only a few genes showing poor prediction scores (b) Relationship between average rank of gene expression and its prediction correlation, suggests that the top expressed values are more difficult to predict compared to lower values (c) Pathways enriched in the top 5% best predicted genes and (d) pathway enrichment of 5% genes most difficult to predict.

#### 3.5.2 Performance table of each gene

Next we try to identify if a pattern exists between the average expression of a gene and its predictability in our model. This analysis is prompted by the observation that regression values are generally better estimated for smaller values due to the nature of activation and objective functions. From Figure 4(b) we do observe that the high expression values are relatively less well predicted than the lower expressed cases. Nonetheless, even in the worst cases, we get sufficiently good estimates of masked gene expression values, giving confidence in the usability of the model.

#### 3.5.3 Functional analysis of top and worst performing genes

Finally, we try to explore if there are specific functional patterns in the best and worst predicted genes. This analysis is relevant because in CMAP project certain best inferred genes were selected which could be best predicted from a small subset of 100 genes. In this case, all the CMAP genes, including best inferred in that work are predictor genes. Here we perform the analysis for the genes absent in GPL96 and GPL571, which are actually not considered at all in CMAP project. Figure 4(c-d) show the functional enrichment of top and bottom 5% genes sorted by their performance (full list of these genes is provided in Supplementary Table ST5). Such a question is relevant because estimating the expression values for some genes from the others is possible due to network associations between them. Gene sets which represent better connectivity on these networks are therefore likely to be better predicted. Figure 4(c-d) shows that the best predicted genes are associated to mitochondrial function. This has support from literature as studies on genomic interactions show that nuclear genes sensitive to mitochondrial variation often sit within broader cellular processes (such as metabolism and muscle function) and coordinate via complex signaling pathways like the unfolded protein response (Chen et al., 2014). On the other hand the worst predicted genes were enriched in biological functions associated to membranes and high specialized functions such as fertility and reproduction, which are intuitively not well connected with other genes, which may contribute to their lower predictability from the measured predictor-gene set. However, even among the approximately 330 genes comprising the lowest-performing 5% of the target set, the mean Pearson correlation was still ∼0.683, suggesting that despite the lower performance of this subset majority of target genes could be predicted very well.

### 3.6 Comparison with related methods and tools

Table 3 highlights the differences between the available integrated resources on large scale microarray data and shows the gaps the MPGEM is trying to bridge.

**Table 3:** Comparison of MPGEM capabilities with similar resources on microarray data.

| Tool/ Feature | Last updated | Frozen/ fixed reference normalization | Subset normalization | Cross-platform data | Transcriptome completion (imputation) |
| --- | --- | --- | --- | --- | --- |
| DGEX | 2016 | No | No | No | Within platform; fixed inputs/ target sets |
| fRMA | 2010 | Yes | No | Yes | No |
| Shambhala-2 | 2022 | Yes | No | No | No |
| Refine.bio | 2026 | Yes | No | Yes | No |
| GEMMA | 2021 | Yes | No | Yes | No |
| MPGEM | Current work | Yes | Yes | Yes | Yes; scalable |

### 3.7 MPGEM-engine performance comparison

Because transcriptome completion of platform-specific, unmeasured genes is not directly addressed by most existing methods, we benchmarked the MPGEM Engine against two simple prediction baselines: gene-wise mean imputation and K-nearest neighbors (KNN). For gene-wise mean imputation, each masked target value was replaced by the mean expression of that gene in the available data. For KNN, the masked expression value was estimated from the corresponding values in the K most similar samples. KNN provides a useful non-parametric reference, but its inference requires continued access to the reference dataset.

In the controlled benchmark, MPGEM achieved slightly higher overall performance than KNN. The mean sample-wise Pearson and Spearman correlations were 0.9436 and 0.9393, respectively, compared with 0.9413 and 0.9378 for KNN. The advantage was more pronounced among the most difficult 5% of samples, for which MPGEM achieved Pearson and Spearman correlations of 0.6733 and 0.6491, compared with 0.6369 and 0.6096 for KNN. MPGEM also showed higher gene-wise correlations than KNN, with Pearson and Spearman values of 0.8301 and 0.8249, compared with 0.8228 and 0.8109, respectively (Table 4a).

**Table 4:** Comparison of MPGEM engine performance with Mean value assignment and KNN-based predictions (a) Pearson’s and Spearman’s correlation coefficients for each model in sample-wise, overall gene-wise and 5% tail gene-wise from each method and p-values from a paired t-test for comparison of mean performance levels between MPGEM engine and KNN based prediction performances (b) Deployment load for usage of new samples in each method.

| Metric | Mean Imputation | KNN (K=5) | MPGEM Engine | p-value (KNN vs MPGEM engine) |
| --- | --- | --- | --- | --- |
| <u>Sample-wise (overall)</u><br>Pearson's Correlation Coefficient<br>Spearman's Correlation Coefficient | 0.7952<br>0.7921 | 0.9413<br>0.9378 | 0.9436<br>0.9393 | 2.45e-70<br>1.79e-27 |
| <u>Sample-wise (tail 5%)</u><br>Pearson's Correlation Coefficient<br>Spearman's Correlation Coefficient | 0.2155<br>0.2287 | 0.6369<br>0.6096 | 0.6733<br>0.6491 | 1.52e-115<br>3.45e-142 |
| <u>Gene-wise (overall)</u><br>Pearson's Correlation Coefficient<br>Spearman's Correlation Coefficient | NA | 0.8228<br>0.8109 | 0.8301<br>0.8249 | 4.87e-231<br><1.0e-300 |
| <u>Gene-wise (tail 5%)</u><br>Pearson's Correlation Coefficient<br>Spearman's Correlation Coefficient | NA | 0.6731<br>0.6298 | 0.6825<br>0.6537 | 3.24e-39<br>8.76e-97 |

(b)
| Property | MPGEM Engine | KNN (K=5) |
| --- | --- | --- |
| Requires training/reference samples at inference | No | Yes |
| Portable trained model | Yes | No* |
| Prediction for large numbers of local samples | Highly suitable | Reference -data dependent |
| Storage after model training | Model parameters only | Reference dataset/index |
| Architecture | Parametric MLP | Non-parametric |

\* KNN requires access to the full reference dataset (or a search index over it) at inference time, rather than a self-contained portable model.

These results indicate that the MPGEM Engine provides performance comparable to, and modestly better than, KNN overall, with a larger advantage for difficult predictions. In addition, the trained MPGEM Engine is a portable parametric model that can be distributed and applied to new samples without retaining the reference samples required by KNN. This makes MPGEM particularly suitable for local deployment on large numbers of samples. The comparison of deployment requirements is summarized in Table 4b.

Although the MPGEM Engine outperformed the evaluated baselines in this benchmark, alternative prediction models may provide further improvements and can be incorporated into the MPGEM framework because the harmonization and prediction components are modular.

### 3.7 Software availability

The pipeline is distributed as an open-source Python package, mpgem, at https://github.com/SciWhylab/MPGEM_TOOL, released under the MIT License. It provides a command-line and Python interface that applies the reference normalization (2.3) and the MPGEM-engine imputation (2.4) to a user-supplied expression matrix. It accepts probe-level or gene-level input; probe-level input is collapsed by the max-per-gene rule (2.2.1), and genes absent from the reference are dropped and logged. The GPL570 reference statistics and the trained engine are bundled, so no external download is required. The MPGEM matrix (207,135 samples × 19,320 genes) is also available through a query interface at http://sciwhylab.jnu.ac.in/servers/mpgem/.

RSQD can also be applied to any expression profile whose genes are a subset of the GPL570 using the on-demand RSQD procedure. Imputation is available when the input supplies exactly the 12,712-gene predictor set shared by GPL571 and GPL96: the normalized profile is passed to the trained MPGEM-engine, which predicts the 6,608 target genes and returns a complete 19,320-gene profile. Profiles with a different gene set are normalized but not imputed, since the engine requires its full 12,712-gene input. An RSQD-lite is provided for portable usage in which subset rank estimates are carried out by pre-averaged expression values of all genes in the GPL570 data and then mapping the ranks of a subset onto it without real-time averages of that subset. Our benchmark shows that such normalization produces the original RSQD described above with very little error (See Supplementary Figure SF1). This portable version is made available in the GitHub repository stated above.

## 4 Discussion and Conclusion

The rapid expansion of public transcriptomic repositories has created unprecedented opportunities for data-driven biological discovery. However, the full potential of these resources has remained largely unrealized because gene expression datasets generated over the past two decades are fragmented across multiple microarray platforms that differ in probe composition, gene coverage, measurement scales, and preprocessing procedures. In this study, we addressed two of the principal obstacles to large-scale integration of legacy microarray data: robust cross-platform harmonization and incomplete transcriptome coverage. We developed a harmonization framework based on the Reference Quantile Distribution (RQD) and its generalized implementation, the Reference Subset Quantile Distribution (RSQD), to transform heterogeneous expression profiles onto a common quantitative scale. Building upon this harmonized representation, we developed the MPGEM Engine, a multilayer perceptron (MLP)-based model that predicts expression values for genes absent from lower-density microarray platforms. Together, these methods enabled the construction of the Multi- Platform Gene Expression Matrix (MPGEM), a unified resource containing harmonized and expanded human gene expression profiles suitable for large-scale computational analyses.

The most significant methodological advance of this work is the extrapolation of expression values for genes that were never experimentally measured on the source microarray platform. This concept is not without precedent. The Connectivity Map (CMAP) (Lamb et al., 2006) and the subsequent LINCS L1000 project (Chen et al., 2016; Subramanian et al., 2017) demonstrated that a carefully selected subset of landmark genes contains sufficient information to reconstruct the expression of thousands of additional genes. In the L1000 framework, expression values of 978 landmark genes were used to infer approximately 11,300 target genes, providing a practical solution for large-scale perturbational profiling. Rather than adopting this implementation directly, we generalized its underlying principle to heterogeneous public microarray datasets. Instead of relying on a predefined landmark panel, our approach exploits all genes measured on each platform as predictive features while treating genes absent from other platforms as prediction targets. This strategy maximizes the biological information available within each experiment and naturally accommodates platforms with widely differing transcriptome coverage.

The success of this strategy is supported by fundamental principles of transcriptional biology. Gene expression is governed by coordinated regulatory networks, signaling pathways, transcription-factor activities, and cell-state-specific programs that generate stable patterns of co-expression across genes. These biological relationships are properties of the underlying cellular system rather than of the experimental platform used to measure them. Although platform differences introduce substantial technical variation, many gene-expression relationships reflect underlying biological programs that can remain sufficiently conserved across datasets to support cross-platform prediction. Consequently, once platform-specific technical variation has been minimized through harmonization, predictor-target relationships learned from one platform become transferable to another. Our RQD/RSQD framework provides precisely this normalization step, transforming heterogeneous datasets onto a common reference distribution and thereby enabling reliable cross-platform gene expression imputation. The high prediction accuracy achieved for masked genes, with average Pearson correlations approaching 0.944, demonstrates that these conserved transcriptional relationships can be effectively exploited to reconstruct substantial portions of the transcriptome.

Compared with existing approaches, MPGEM extends both the scope and applicability of transcriptome reconstruction. Previous inference frameworks, including CMAP and L1000, aimed at predicting drug responses from a minimal gene set and were developed using curated datasets generated from a single microarray platform and therefore operated within a fixed transcriptomic feature space. In contrast, MPGEM is designed with a broader objective explicitly for cross-platform integration. Using GPL570 as the reference transcriptome, the predictor space consists of all genes measured on each source platform rather than a predefined landmark set. Across the three most widely used human microarray platforms, this corresponds to 12,712 core predictor genes, including the 978 L1000 landmark genes, 9,196 CMAP BING genes, and an additional 2,538 genes shared across the three platforms. The remaining 6,608 genes unique to GPL570 constitute the prediction targets, enabling reconstruction of the complete 19,320-gene transcriptome represented on the reference platform. This substantially broader predictor space likely contributes to the strong predictive performance observed in this study while simultaneously preserving a much larger fraction of the original biological information than conventional cross-platform analyses that retain only shared genes.

The implications of this work extend beyond transcriptome harmonization. Large-scale AI models require consistent feature representations across all samples, an assumption that has traditionally limited the use of heterogeneous public gene expression repositories. Differences in gene coverage and measurement scales have forced investigators either to discard large numbers of genes or to restrict analyses to individual platforms, thereby limiting statistical power and reducing biological diversity. By jointly harmonizing expression values and reconstructing missing genes, MPGEM transforms fragmented public microarray collections into a unified transcriptomic matrix that is suitable for meta-analysis, biomarker discovery, disease classification, drug-response prediction, systems biology, and the development of foundation-scale AI models. The resource also provides a standardized framework for incorporating newly generated microarray datasets, allowing future studies to be integrated into an existing harmonized transcriptomic landscape with minimal additional preprocessing.

MPGEM is not dramatically more accurate than KNN overall; it is slightly more accurate and considerably better on difficult cases, while being deployable as a self-contained trained model. Several limitations should nevertheless be acknowledged. First, the accuracy of transcriptome reconstruction depends on the conservation of gene co-expression relationships across biological contexts. Although these relationships are generally robust across diverse tissues and experimental conditions, platform-independent prediction may become less accurate for highly context-specific or unique regulatory programs or genes with weak correlation to the remaining transcriptome. Second, the current implementation focuses on the three most widely used human microarray platforms, using GPL570 as the reference transcriptome. Extending the framework to additional platforms, other species, and transcriptomic technologies will further enhance the breadth and utility of the resource. Finally, while the present study demonstrates excellent reconstruction accuracy using a classical multilayer perceptron, emerging deep-learning architectures, including graph neural networks, transformer-based models, and multimodal foundation models, may further improve transcriptome completion by exploiting higher-order biological relationships.

These limitations also define promising directions for future research. The availability of a harmonized, gene-complete transcriptomic resource opens opportunities for developing large-scale self-supervised foundation models trained directly on public gene expression repositories. Such models could learn transferable representations of cellular states and biological perturbations that support a wide range of downstream applications, including disease diagnosis, drug repurposing, pathway inference, causal network reconstruction, and multimodal integration with genomic, proteomic, spatial transcriptomic, and clinical datasets. Because the RQD/RSQD framework is independent of any specific prediction model, future generations of transcriptome inference algorithms can readily be incorporated into the MPGEM pipeline without requiring changes to the underlying harmonization strategy.

In conclusion, MPGEM establishes a unified computational framework for integrating heterogeneous legacy microarray datasets through robust cross-platform harmonization and AI-driven transcriptome completion. By combining a novel normalization methodology with accurate prediction of genes absent from individual platforms, the framework overcomes two major barriers that have historically limited the large-scale reuse of public gene expression repositories. The resulting resource preserves the scientific value of more than 200,000 legacy transcriptomic profiles while transforming them into a coherent, analysis-ready knowledge base for modern computational biology. We anticipate that MPGEM will accelerate transcriptomic meta-analysis, biomarker discovery, and AI-driven functional genomics, while providing a scalable foundation for seamless integration of legacy microarray data with RNA sequencing and future transcriptomic technologies.

## 6 Author Contributions

SA Conceptualized and coordinated the project, including final MS editing and study design. SG and AKV curated the data, developed the MPGEM engine and performed prediction experiments. SG innovated the normalization and harmonization strategies including RQSD, mined literature and prepared the current manuscript. SJ contributed in software development and visualization, preparing GitHub repositories and performing baseline experiments for benchmarking. All authors read and consented to the final version of the MS.

## 8 Acknowledgments

Authors would like to acknowledge help from Anima Kujur in running codes and Arfa Jabin for conducting tests (not included in this manuscript) in the early stages of this work.

## 10. Supplementary Material

All supplementary tables and results are available from a single file supplementary-all.pdf

## 11. Data Availability Statement

The MPGEM resource together with normalization and imputation pipelines is distributed as an open-source Python package, MPGEM, at https://github.com/SciWhylab/MPGEM_TOOL, released under the MIT License.

---

**Supplementary Table ST1:**
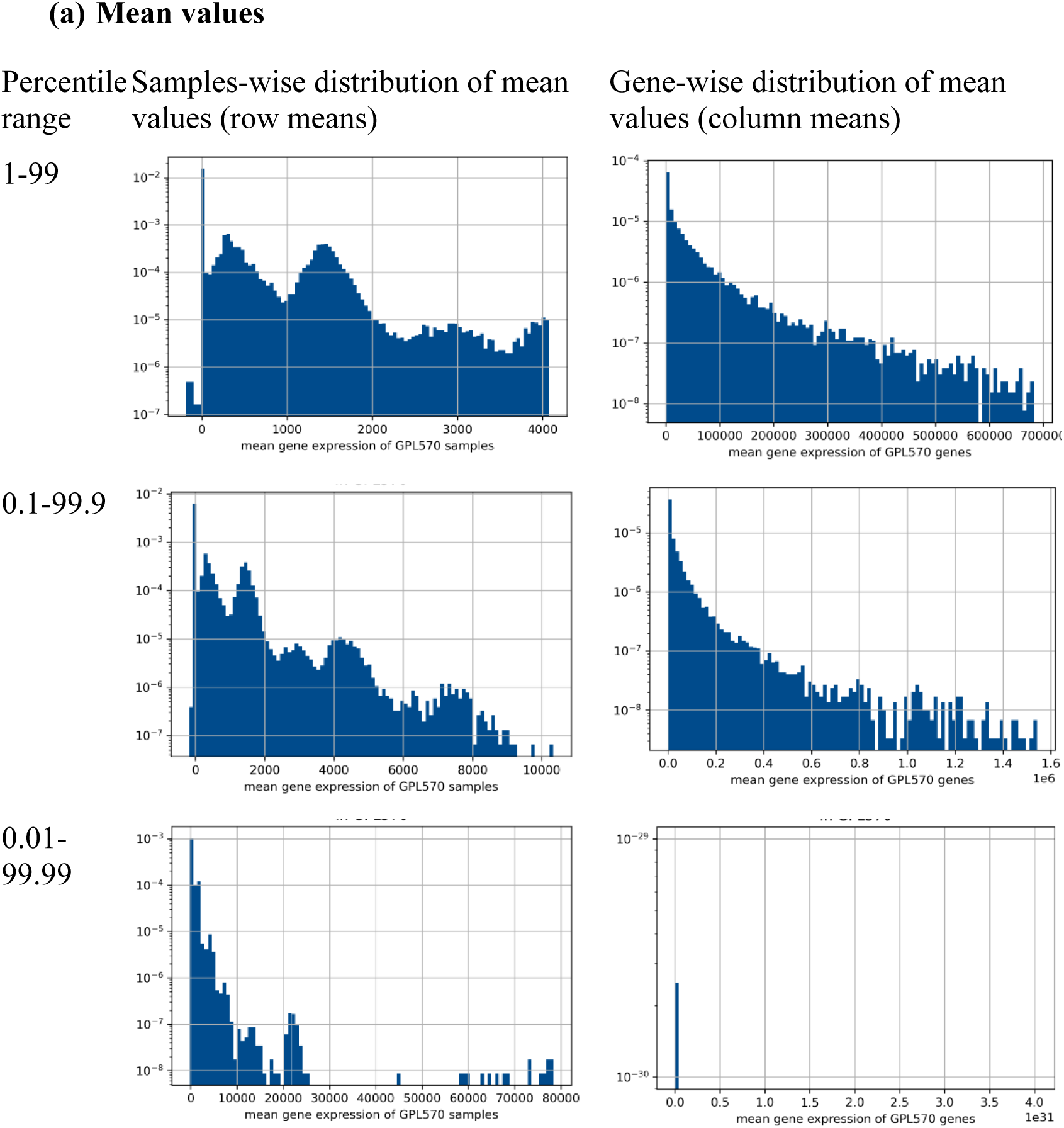
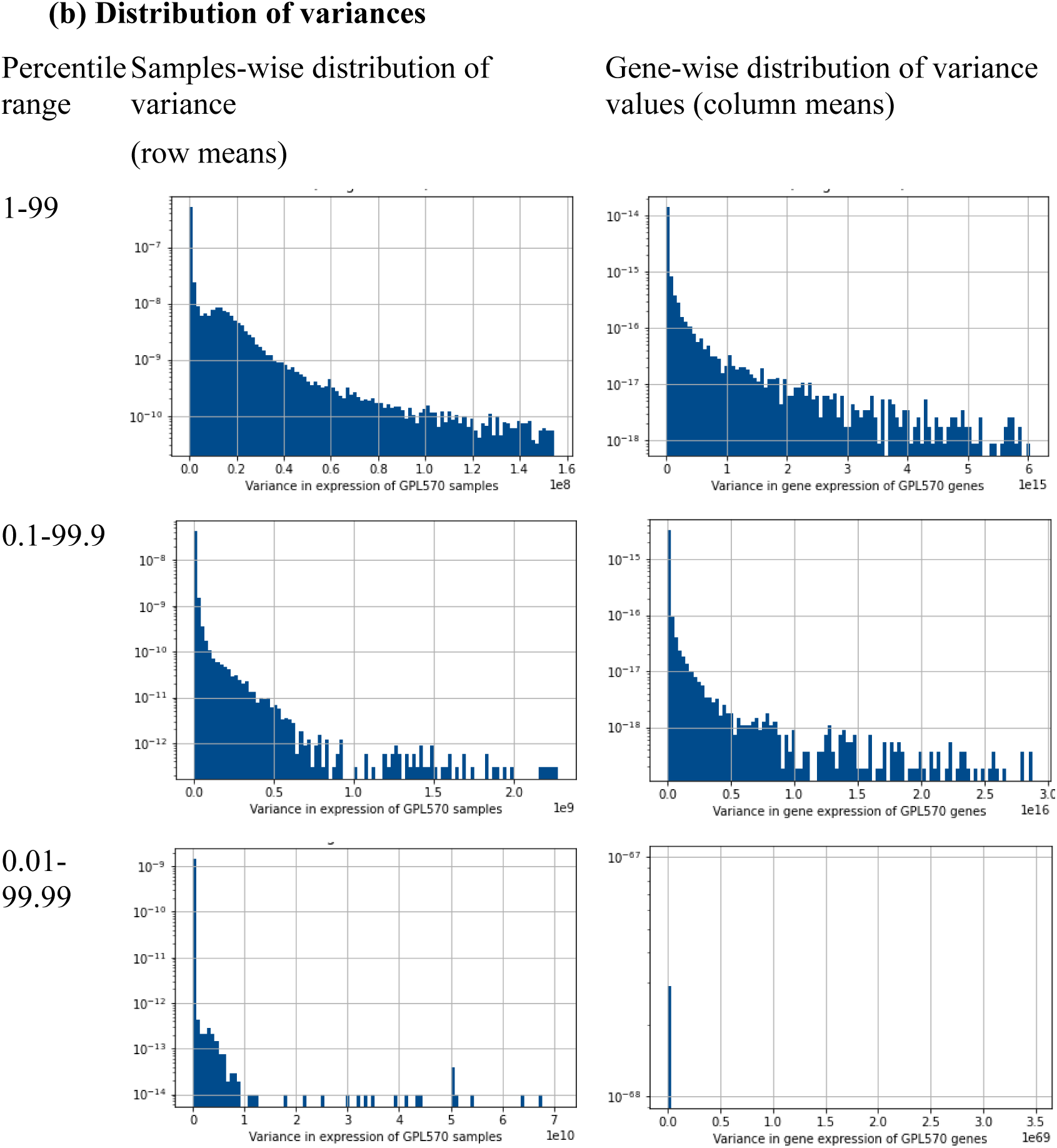
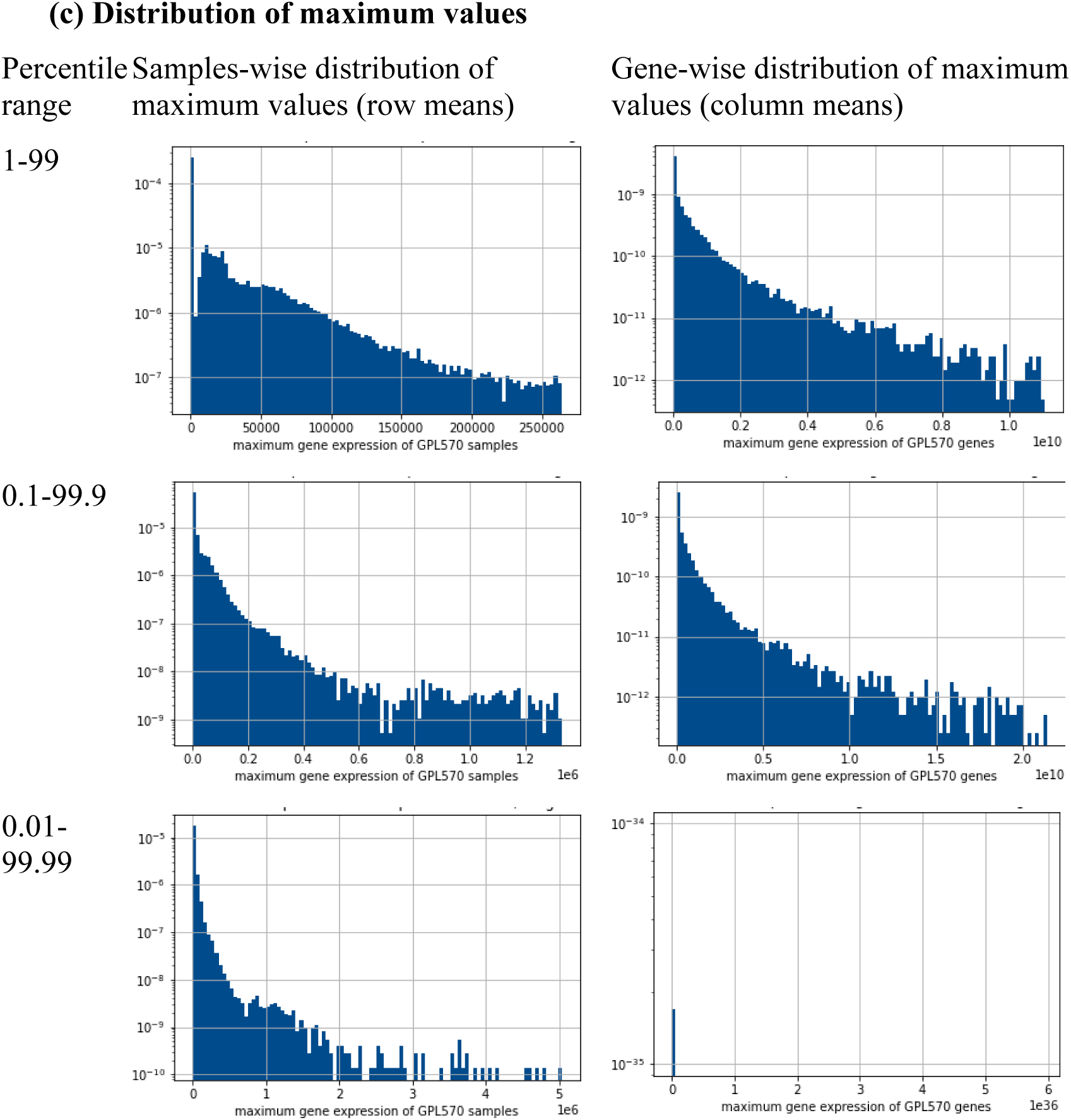
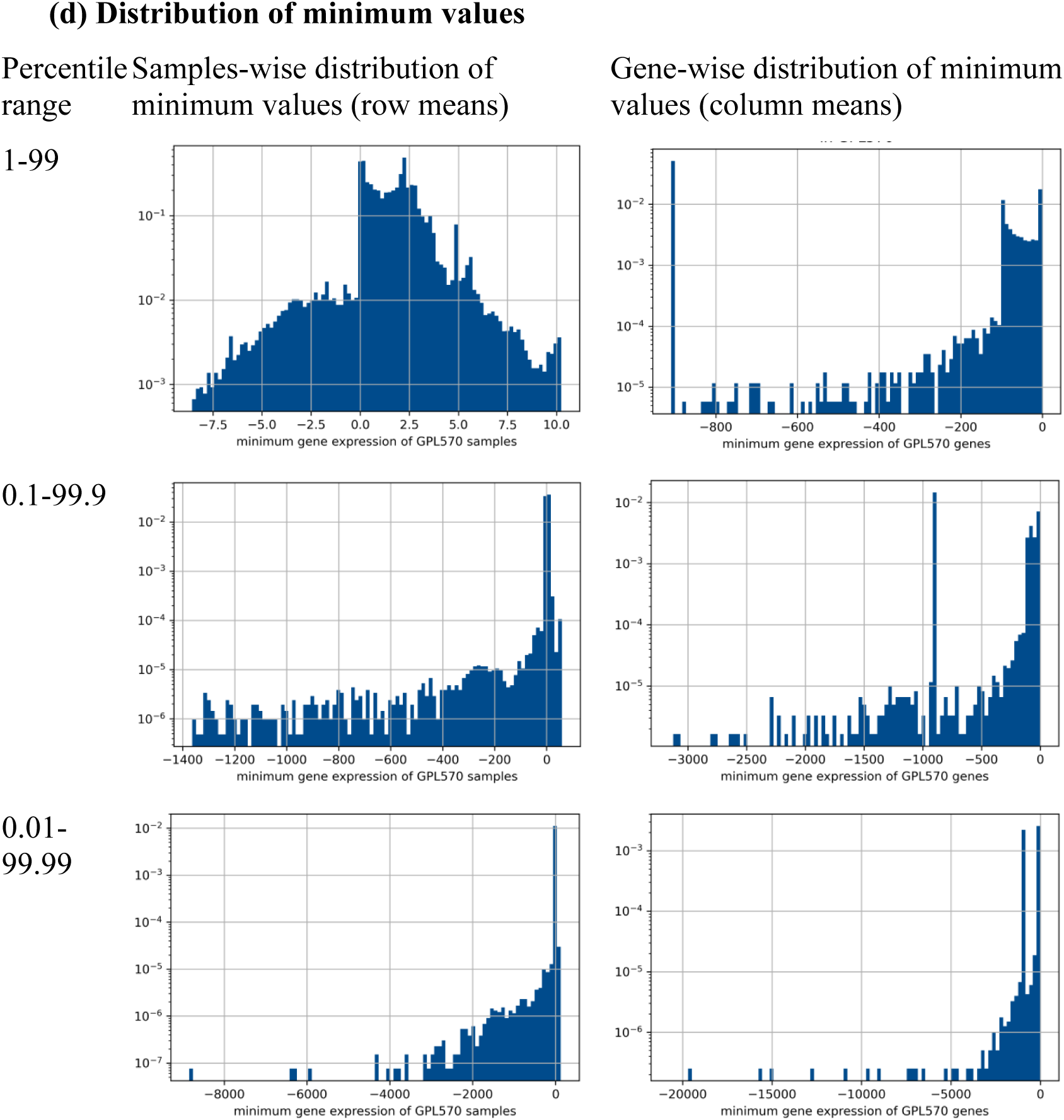
Distribution of mean and variance values in pre-MPGEM: GPL570. Distribution of mean and variance values of sample-wise and gene-wise raw expression values in pre-MPGEM: GPL570 after trimming values outside of their selected percentile ranges (to improve visualization) (a) distribution of mean values (b) distribution of variance (c) distribution of maximum values (d) distribution of minimum values

**Supplementary Table ST2:**
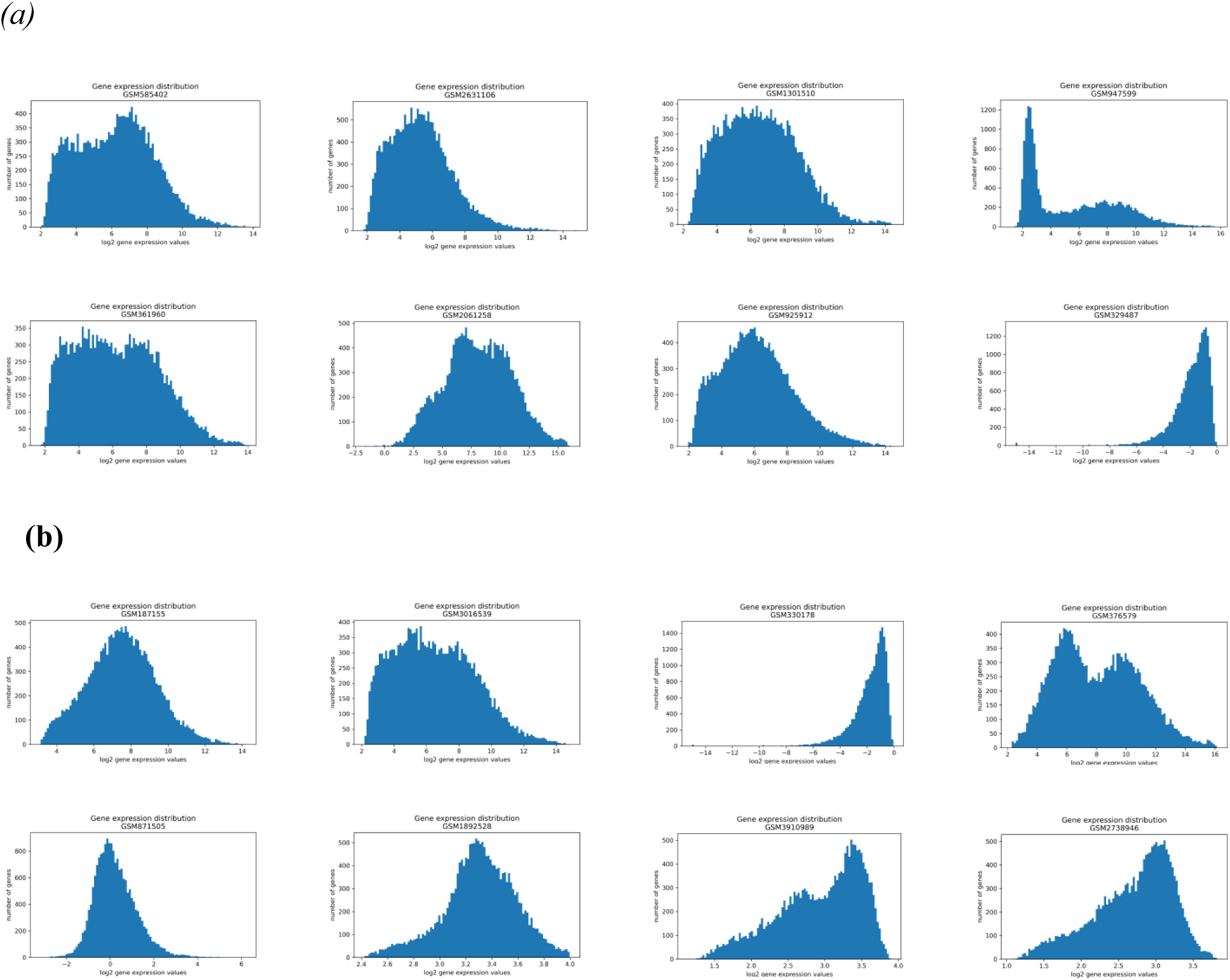
Typical sample-wise gene expression distribution patterns in pre-MPGEM:GPL570 (a) Randomly selected samples (b) Samples selected by Keyword of log2-scaled quantile normalized samples.

**Supplementary Table ST3:**
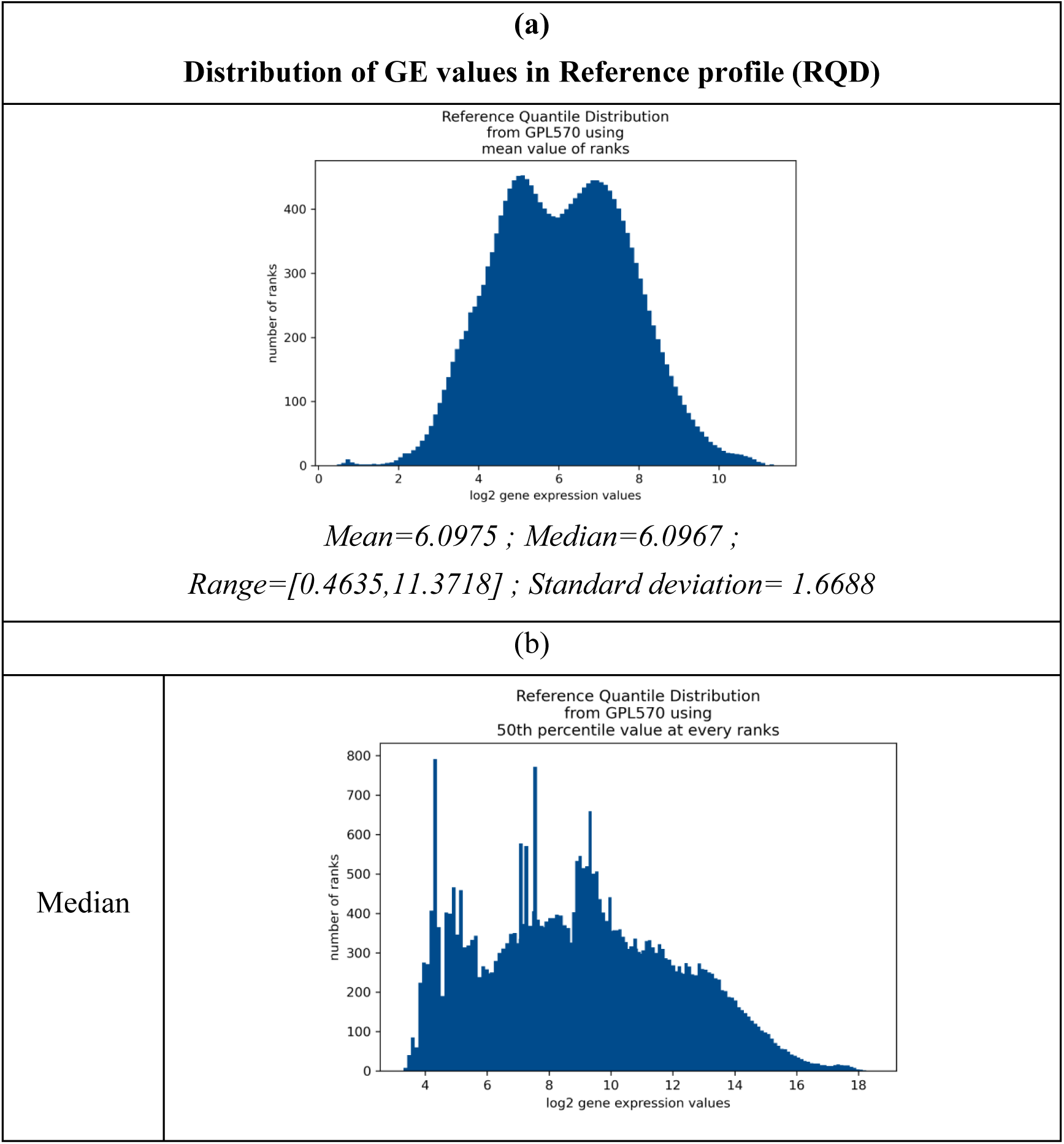
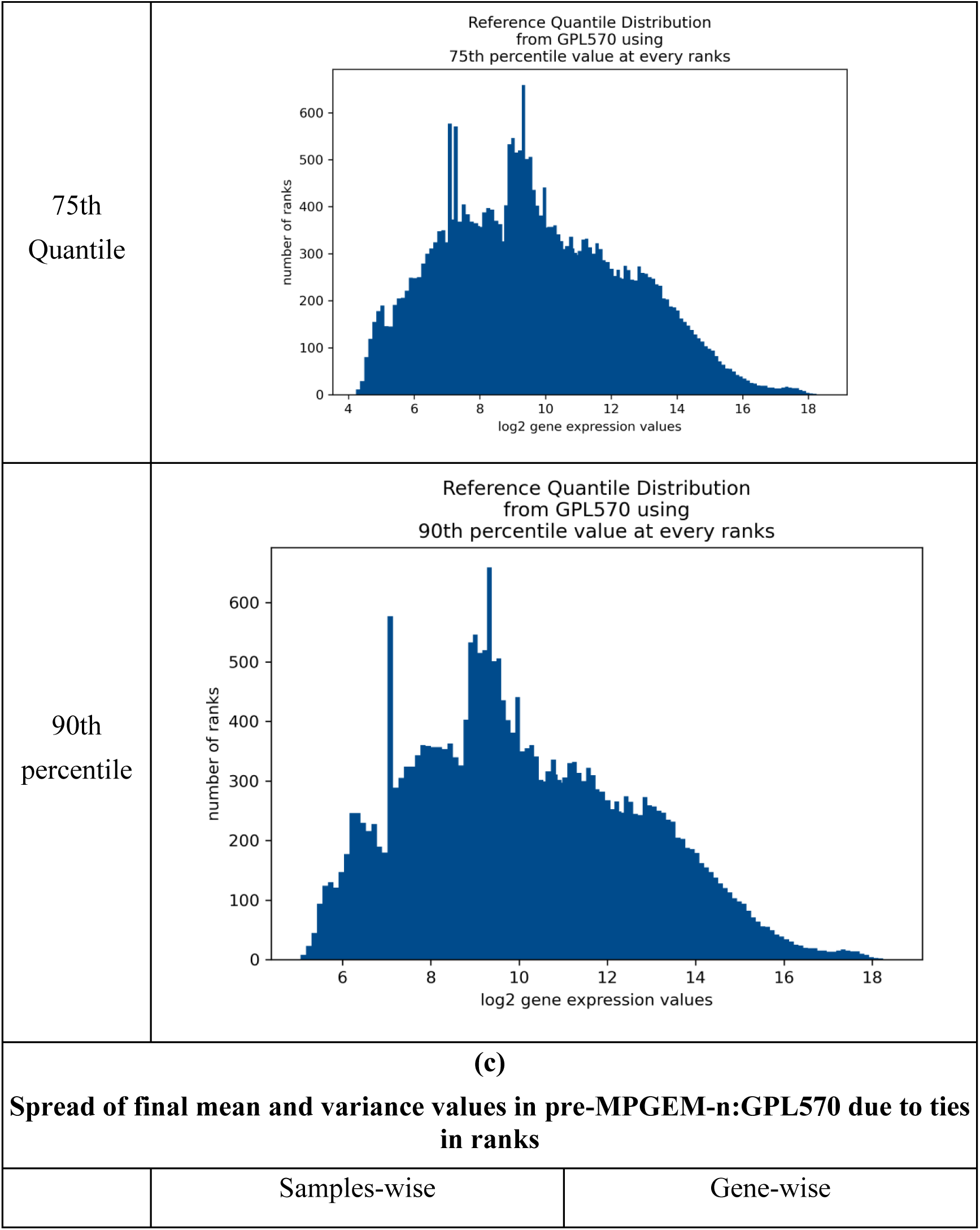
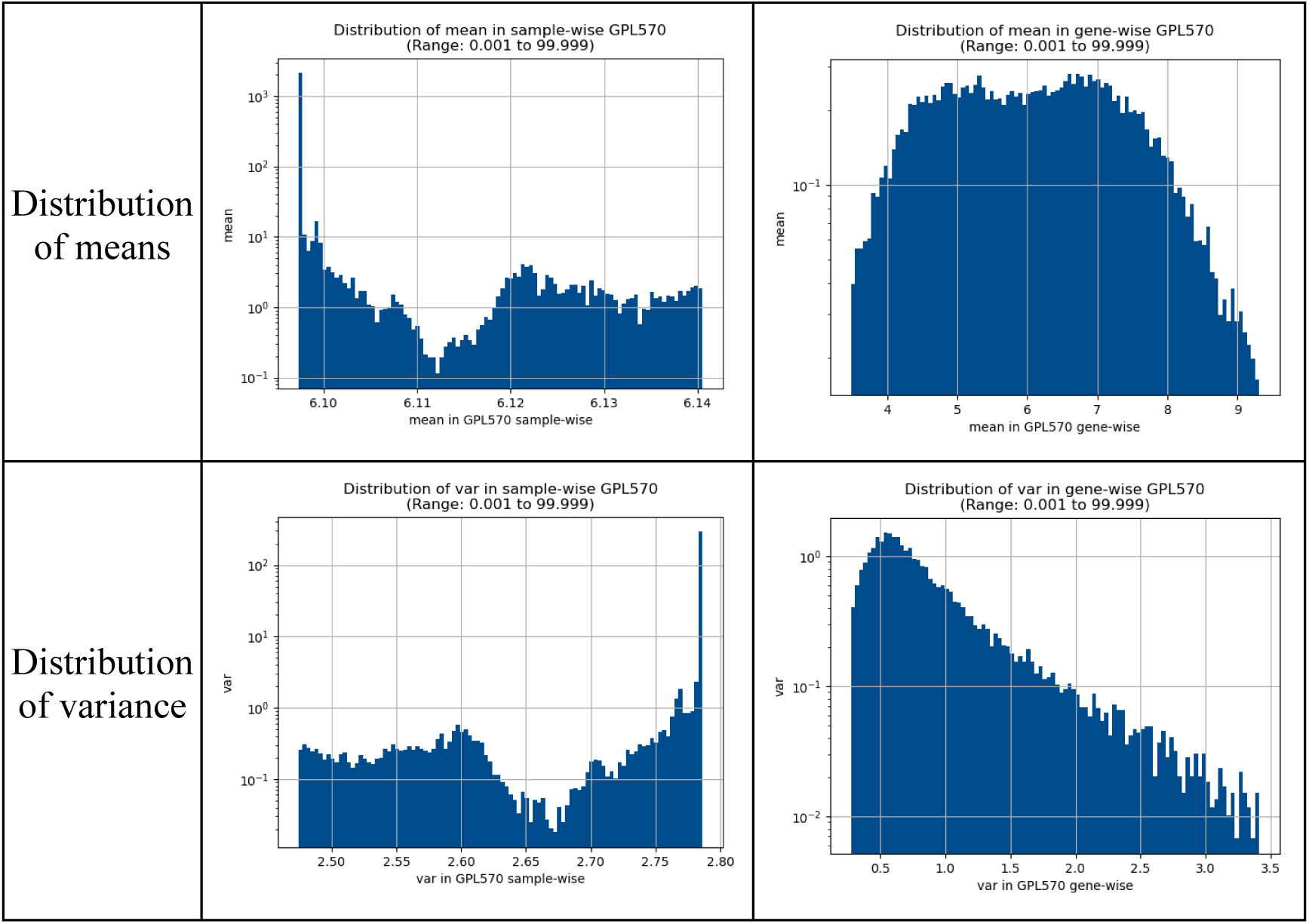
Nature of reference distribution proposed for universal harmonization of gene expression data. (a) Distribution of GE values in Reference profile (RQD) (b) Candidate RQDs based on percentiles of equally-ranked gene expression values across pre-MPGEM:GPL570 (c) Spread of final mean and variance values in pre-MPGEM-n:GPL570 due to ties in ranks

**Supplementary Table ST4:**
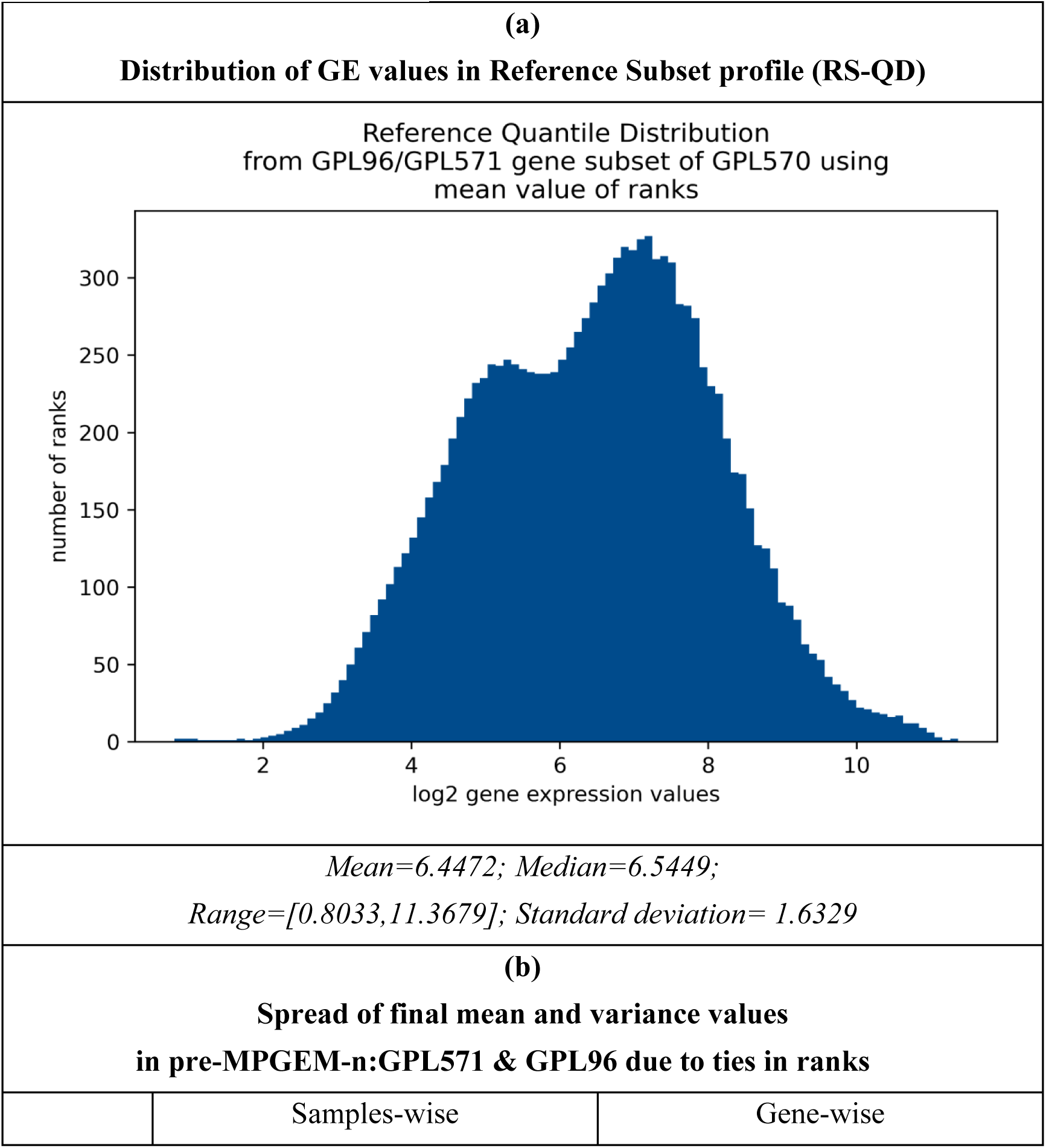
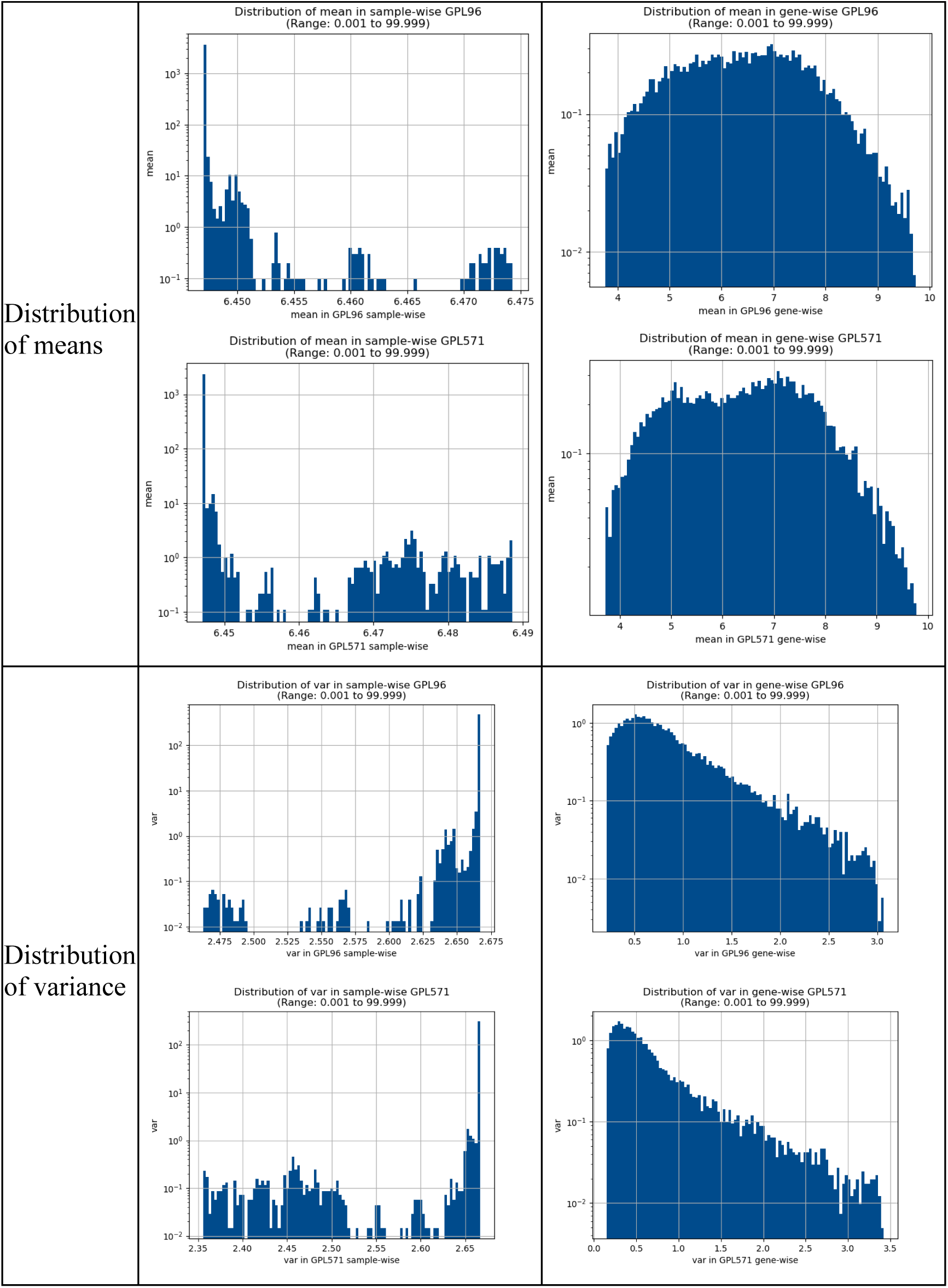
Quantile distribution of GPL96 and GPL571 based in subset referenced RQD.

**Supplementary table ST5:**
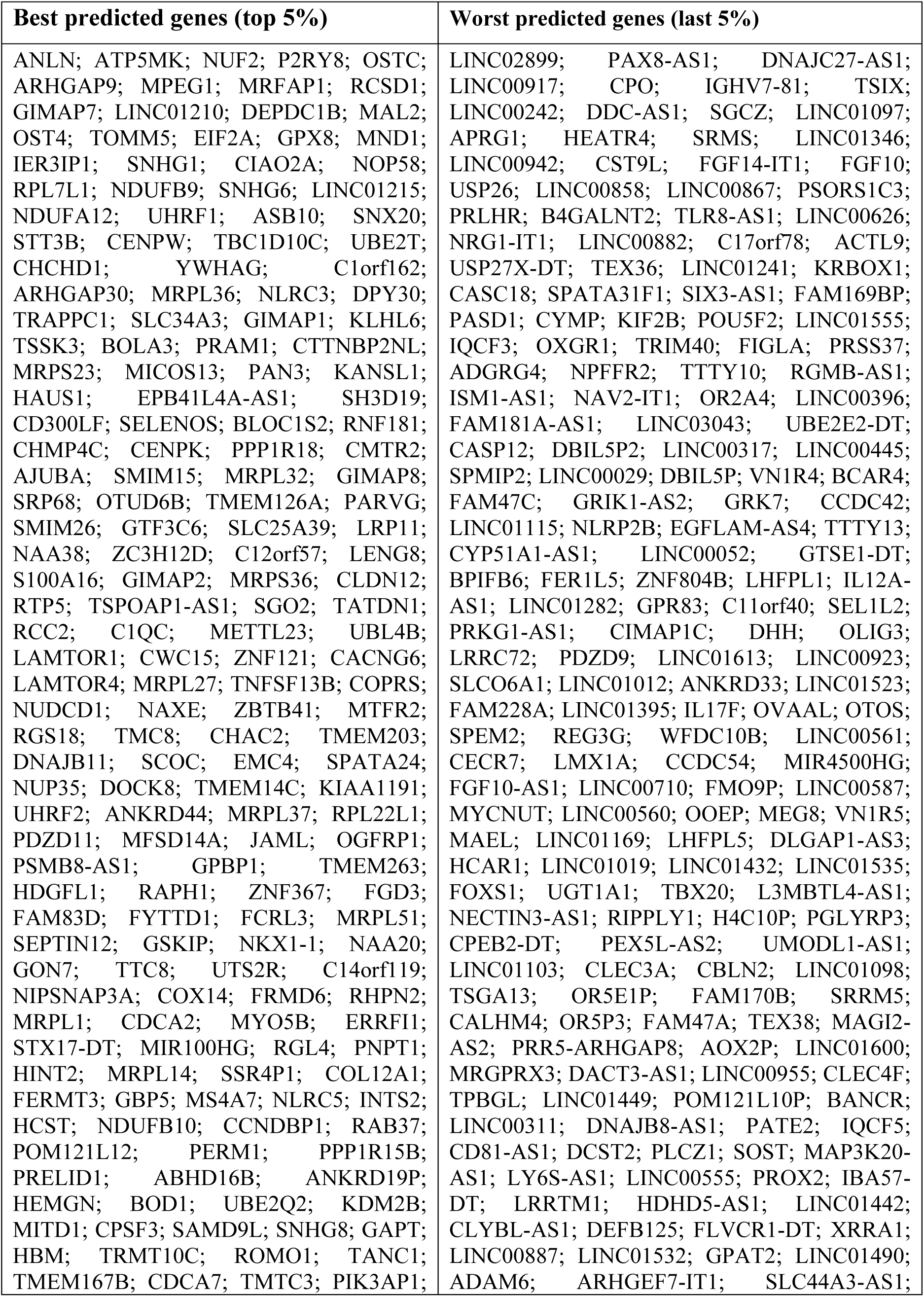
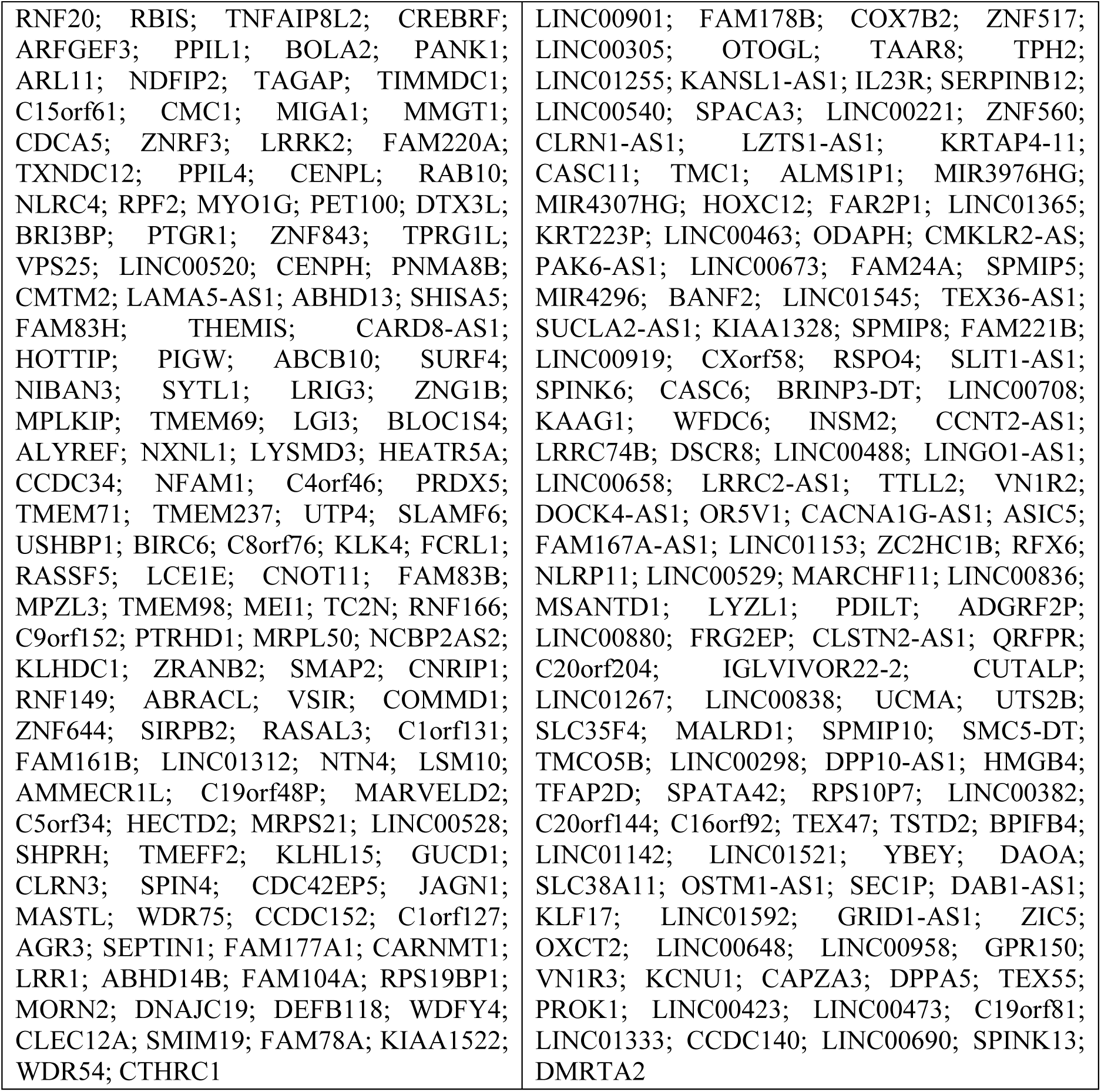
Best and worst predicted genes in MPGEM engine.

**Supplementary Figure SF1:**
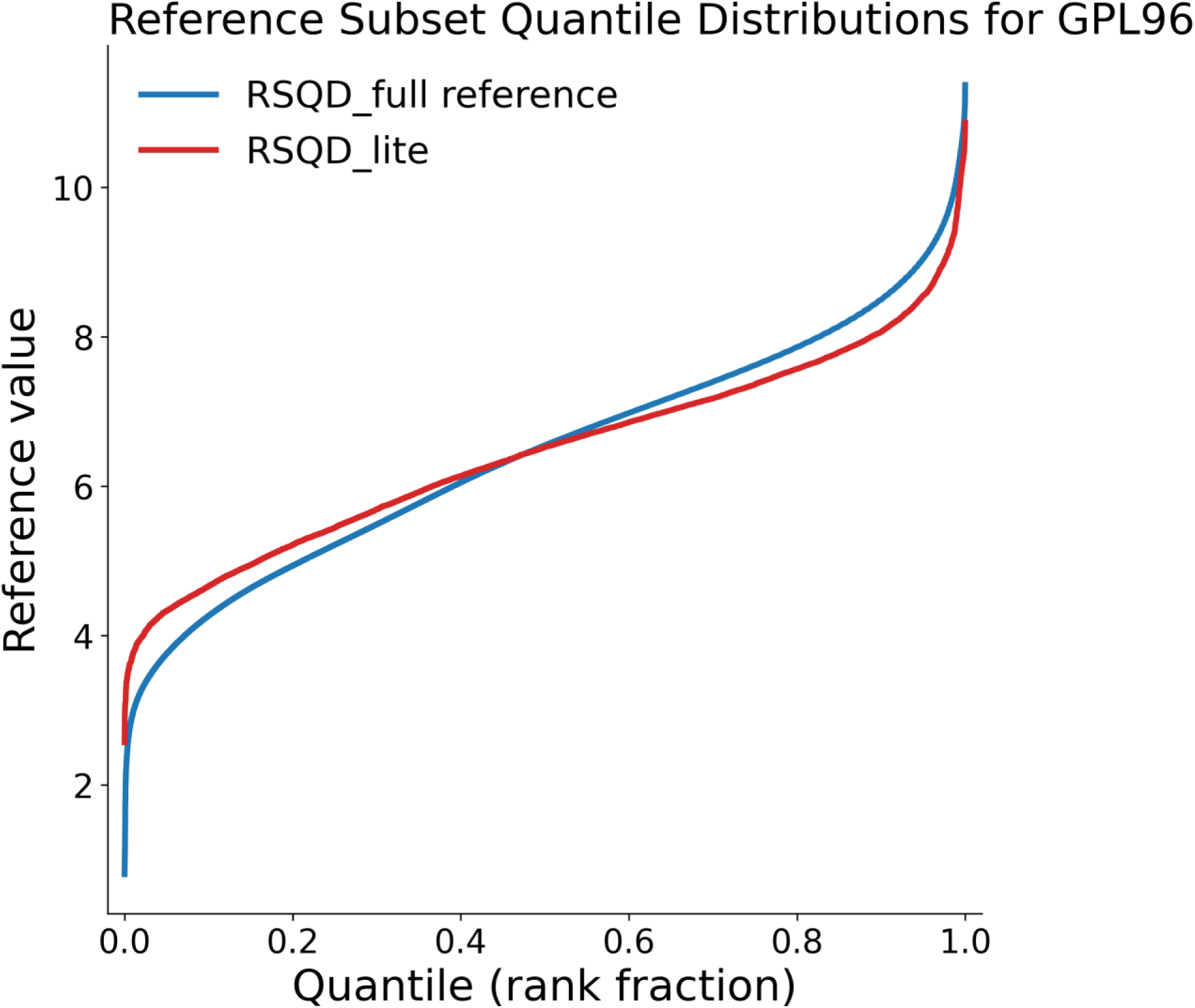
This figure shows a comparison between the the normalized values of gene expression data from GPL96 obtained by full RSQD compared to its lite version with pre-computed mean values. There is a good agreement between the two sets of values and most divergence happens in the extreme rank positions, allowing for portability of method without much loss.

## References

Abadi, M. and others (2015). TensorFlow: Large-Scale Machine Learning on Heterogeneous Systems. Available at: https://www.tensorflow.org/

Allen, J. D., Wang, S., Chen, M., Girard, L., Minna, J. D., Xie, Y., et al. (2012). Probe mapping across multiple microarray platforms. Briefings in Bioinformatics 13, 547–554. doi: 10.1093/bib/bbr076

Athar, A., Füllgrabe, A., George, N., Iqbal, H., Huerta, L., Ali, A., et al. (2019). ArrayExpress update – from bulk to single-cell expression data. Nucleic Acids Research 47, D711–D715. doi: 10.1093/nar/gky964

Bolstad, B. M., Irizarry, R. A., Åstrand, M., and Speed, T. P. (2003). A comparison of normalization methods for high densityoligonucleotide array data based on variance and bias. Bioinformatics 19, 185–193. doi: 10.1093/bioinformatics/19.2.185

Borisov, N., Shabalina, I., Tkachev, V., Sorokin, M., Garazha, A., Pulin, A., et al. (2019). Shambhala: a platform-agnostic data harmonizer for gene expression data. BMC Bioinformatics 20, 66. doi: 10.1186/s12859-019-2641-8

Borisov, N., Sorokin, M., Zolotovskaya, M., Borisov, C., and Buzdin, A. (2022). Shambhala-2: A Protocol for Uniformly Shaped Harmonization of Gene Expression Profiles of Various Formats. Current Protocols 2, e444. doi: 10.1002/cpz1.444

Chen, C., Hyun, T. K., Han, X., Feng, Z., Li, Y., Liu, X., et al. (2014). Coexpression within Integrated Mitochondrial Pathways Reveals Different Networks in Normal and Chemically Treated Transcriptomes. International Journal of Genomics 2014, 1–10. doi: 10.1155/2014/452891

Chen, Y., Li, Y., Narayan, R., Subramanian, A., and Xie, X. (2016). Gene expression inference with deep learning. Bioinformatics 32, 1832–1839. doi: 10.1093/bioinformatics/btw074

Chollet, F. and others (2015). Keras. Available at: https://github.com/keras-team/keras

Clough, E., Barrett, T., Wilhite, S. E., Ledoux, P., Evangelista, C., Kim, I. F., et al. (2024). NCBI GEO: archive for gene expression and epigenomics data sets: 23-year update. Nucleic Acids Research 52, D138–D144. doi: 10.1093/nar/gkad965

Cooper, C. S., Campbell, C., and Jhavar, S. (2007). Mechanisms of Disease: biomarkers and molecular targets from microarray gene expression studies in prostate cancer. Nat Rev Urol 4, 677–687. doi: 10.1038/ncpuro0946

Edgar, R. (2002). Gene Expression Omnibus: NCBI gene expression and hybridization array data repository. Nucleic Acids Research 30, 207–210. doi: 10.1093/nar/30.1.207

Ganter, B., Snyder, R. D., Halbert, D. N., and Lee, M. D. (2006). Toxicogenomics in Drug Discovery and Development: Mechanistic Analysis of Compound/Class-Dependent Effects Using the Drugmatrix^®^ Database. Pharmacogenomics 7, 1025–1044. doi: 10.2217/14622416.7.7.1025

Glorot, X., and Bengio, Y. (2010). Understanding the difficulty of training deep feedforward neural networks., in Proceedings of the Thirteenth International Conference on Artificial Intelligence and Statistics, eds. Y. W. Teh and M. Titterington (Chia Laguna Resort, Sardinia, Italy: PMLR), 249–256. Available at: https://proceedings.mlr.press/v9/glorot10a.html

Golub, T. R., Slonim, D. K., Tamayo, P., Huard, C., Gaasenbeek, M., Mesirov, J. P., et al. (1999). Molecular Classification of Cancer: Class Discovery and Class Prediction by Gene Expression Monitoring. Science 286, 531–537. doi: 10.1126/science.286.5439.531

Greene, C. S., Hu, D., Jones, R. W. W., Liu, S., Mejia, D. S., Patro, R., et al. (2026). refine.bio: a resource of uniformly processed publicly available gene expression datasets. Available at: https://www.refine.bio

Huang, Z., Han, Z., Wang, T., Shao, W., Xiang, S., Salama, P., et al. (2021). TSUNAMI: Translational Bioinformatics Tool Suite for Network Analysis and Mining. *Genomics*, Proteomics & Bioinformatics 19, 1023–1031. doi: 10.1016/j.gpb.2019.05.006

Igarashi, Y., Nakatsu, N., Yamashita, T., Ono, A., Ohno, Y., Urushidani, T., et al. (2015). Open TG-GATEs: a large-scale toxicogenomics database. Nucleic Acids Research 43, D921–D927. doi: 10.1093/nar/gku955

Johnson, W. E., Li, C., and Rabinovic, A. (2007). Adjusting batch effects in microarray expression data using empirical Bayes methods. Biostatistics 8, 118–127. doi: 10.1093/biostatistics/kxj037

Korir, P. K., Geeleher, P., and Seoighe, C. (2015). Seq-ing improved gene expression estimates from microarrays using machine learning. BMC Bioinformatics 16, 286. doi: 10.1186/s12859-015-0712-z

Lamb, J., Crawford, E. D., Peck, D., Modell, J. W., Blat, I. C., Wrobel, M. J., et al. (2006). The Connectivity Map: Using Gene-Expression Signatures to Connect Small Molecules, Genes, and Disease. Science 313, 1929–1935. doi: 10.1126/science.1132939

Lockhart, D. J., Dong, H., Byrne, M. C., Follettie, M. T., Gallo, M. V., Chee, M. S., et al. (1996). Expression monitoring by hybridization to high-density oligonucleotide arrays. Nat Biotechnol 14, 1675–1680. doi: 10.1038/nbt1296-1675

McCall, M. N., Bolstad, B. M., and Irizarry, R. A. (2010). Frozen robust multiarray analysis (fRMA). Biostatistics 11, 242–253. doi: 10.1093/biostatistics/kxp059

Miller, J. A., Cai, C., Langfelder, P., Geschwind, D. H., Kurian, S. M., Salomon, D. R., et al. (2011). Strategies for aggregating gene expression data: The collapseRows R function. BMC Bioinformatics 12, 322. doi: 10.1186/1471-2105-12-322

Ramasamy, A., Mondry, A., Holmes, C. C., and Altman, D. G. (2008). Key Issues in Conducting a Meta-Analysis of Gene Expression Microarray Datasets. PLoS Med 5, e184. doi: 10.1371/journal.pmed.0050184

Rung, J., and Brazma, A. (2013). Reuse of public genome-wide gene expression data. Nat Rev Genet 14, 89–99. doi: 10.1038/nrg3394

Schena, M., Shalon, D., Davis, R. W., and Brown, P. O. (1995). Quantitative Monitoring of Gene Expression Patterns with a Complementary DNA Microarray. Science 270, 467–470. doi: 10.1126/science.270.5235.467

Shabalin, A. A., Tjelmeland, H., Fan, C., Perou, C. M., and Nobel, A. B. (2008). Merging two gene-expression studies via cross-platform normalization. Bioinformatics 24, 1154–1160. doi: 10.1093/bioinformatics/btn083

Stark, J. C., Pipko, N., Liang, Y., Szuto, A., Tsoi, C. T., Dickson, M. A., et al. (2025). Clinical applications of and molecular insights from RNA sequencing in a rare disease cohort. Genome Med 17, 72. doi: 10.1186/s13073-025-01494-w

Su, Z., Fang, H., Hong, H., Shi, L., Zhang, W., Zhang, W., et al. (2014). An investigation of biomarkers derived from legacy microarray data for their utility in the RNA-seq era. Genome Biol 15, 523. doi: 10.1186/s13059-014-0523-y

Subramanian, A., Narayan, R., Corsello, S. M., Peck, D. D., Natoli, T. E., Lu, X., et al. (2017). A Next Generation Connectivity Map: L1000 Platform and the First 1,000,000 Profiles. Cell 171, 1437–1452.e17. doi: 10.1016/j.cell.2017.10.049

Van De Vijver, M. J., He, Y. D., Van ’T Veer, L. J., Dai, H., Hart, A. A. M., Voskuil, D. W., et al. (2002). A Gene-Expression Signature as a Predictor of Survival in Breast Cancer. N Engl J Med 347, 1999–2009. doi: 10.1056/NEJMoa021967

Van ’T Veer, L. J., Dai, H., Van De Vijver, M. J., He, Y. D., Hart, A. A. M., Mao, M., et al. (2002). Gene expression profiling predicts clinical outcome of breast cancer. Nature 415, 530–536. doi: 10.1038/415530a

Walsh, C., Hu, P., Batt, J., and Santos, C. (2015). Microarray Meta-Analysis and Cross-Platform Normalization: Integrative Genomics for Robust Biomarker Discovery. Microarrays 4, 389–406. doi: 10.3390/microarrays4030389

Wang, Z., Gerstein, M., and Snyder, M. (2009). RNA-Seq: a revolutionary tool for transcriptomics. Nat Rev Genet 10, 57–63. doi: 10.1038/nrg2484

Waters, M., Stasiewicz, S., Alex Merrick, B., Tomer, K., Bushel, P., Paules, R., et al. (2007). CEBS Chemical Effects in Biological Systems: a public data repository integrating study design and toxicity data with microarray and proteomics data. Nucleic Acids Research 36, D892–D900. doi: 10.1093/nar/gkm755

Zoubarev, A., Hamer, K. M., Keshav, K. D., McCarthy, E. L., Santos, J. R. C., Van Rossum, T., et al. (2012). Gemma: a resource for the reuse, sharing and meta-analysis of expression profiling data. Bioinformatics 28, 2272–2273. doi: 10.1093/bioinformatics/bts430

